# Cooperative FOXA1–HNF4A binding emerges from motif spacing and nucleosome architecture

**DOI:** 10.64898/2026.05.27.728252

**Authors:** Patrick D. Wilson, Michael J. Buck

## Abstract

The pioneer factor hypothesis posits that specialized transcription factors access nucleosomal DNA to enable binding of secondary factors, implying a hierarchical mechanism of chromatin opening. However, recent evidence indicates that cooperative binding between pioneer and non-pioneer factors can occur in a context-dependent manner. Here we define the sequence and chromatin features that specify cooperative binding between FOXA1 and HNF4A. Using dual-induction of FOXA1 and HNF4A in naïve cells, which lack endogenous expression of either factor, we identify a class of genomic sites that require both factors for binding and reside in initially inaccessible chromatin. A dual-head convolutional neural network finds no novel composite motif at these sites, instead implicating motif arrangement as the distinguishing feature. Cooperative loci are enriched for FOXA1–HNF4A motif pairs spaced 15–60 bp apart, a soft-syntax grammar consistent with nucleosome-mediated cooperativity. This spacing signature recurs at endogenously co-bound sites in HepG2 cells and primary fetal hepatocytes. Using a multiplexed in vitro nucleosome-binding assay (Pioneer-seq), we show that FOXA1 binds nucleosomal DNA more strongly than HNF4A across positions, and that cooperative binding strength on natural nucleosome sequences is predicted primarily by the position of the FOXA1 motif relative to the nucleosome dyad. Together, these results support a model in which positioned nucleosomes act as scaffolds that facilitate cooperative transcription factor binding through spatial motif organization, rather than a barrier one factor clears for the other.

## Introduction

The pioneer factor hypothesis holds that a specialized class of transcription factors can engage nucleosomal DNA in compacted chromatin, establishing accessible states that enable subsequent binding by other regulatory proteins. The foundational evidence for this model came from in vitro studies of the forkhead factor HNF3 (now FOXA1) at the albumin enhancer. Cirillo et al. showed that FOXA1’s winged-helix domain structurally resembles linker histone H1 and can bind the lateral surface of nucleosome cores^1^. In a key follow-up experiment, they reconstituted nucleosome arrays containing albumin enhancer sequences, compacted them with linker histone, and tested which transcription factors could still bind. FOXA1 and GATA-4 bound their sites in compacted chromatin and opened the local nucleosomal domain without ATP-dependent remodeling enzymes, while NF-1, C/EBP, and GAL4 could not^2^. These experiments established a qualitative distinction between “pioneer” factors capable of chromatin engagement and “settler” factors that require pre-existing accessibility.

Subsequent work reinforced this hierarchy in the liver gene regulatory network. In adult mouse liver, FOXA proteins continuously maintain enhancer accessibility, nucleosome positioning, and binding of the nuclear receptor HNF4A; ablation of all FOXA genes causes rapid collapse of the hepatic transcriptional program back to pre-hepatic levels^3^. Single-molecule tracking experiments showed that FOXA1 targets DNase-resistant, stable nucleosomes using slow diffusion and long residence times, while HNF4A preferentially targets open, DNase-sensitive chromatin and scans compact chromatin inefficiently^4^. Together, these studies supported a model in which FOXA1 arrives first at closed chromatin, opens it, and HNF4A subsequently binds the accessible site in a sequential, hierarchical relationship.

Recent functional tests have challenged the simplicity of this model, however. Hansen, Loell, and Cohen tested the pioneer factor hypothesis directly by expressing FOXA1 and HNF4A, individually and together, in K562 lymphoblast cells that lack endogenous expression of either factor^5^. Both transcription factors independently bound and opened inaccessible chromatin sites, activated endodermal genes, and pioneered for each other, though FOXA1 required fewer copies of its motif for binding. A subset of target sites required both factors for binding or activation, but the predominant mode of action at these cooperative sites did not conform to the sequential activity predicted by the pioneer factor hypothesis. In a follow-up study using doxycycline titration to quantify binding affinity at thousands of genomic loci, HNF4A displayed stronger pioneer activity than FOXA1, binding inaccessible sites at lower TF concentrations relative to accessible ones^6^. These findings led the authors to propose that pioneer activity is a quantitative, context-dependent property of all transcription factors — governed by TF-DNA binding affinity and motif content — rather than a qualitative trait of a select class.

The Hansen et al. experiments revealed three functionally distinct classes of co-bound sites: those where FOXA1 alone was sufficient for binding (FOXA1-enabled), those where HNF4A alone sufficed (HNF4A-enabled), and cooperative sites where both factors were required. What distinguishes these classes at the DNA sequence level remains unclear. In particular, Hansen and Cohen quantified binding through the aggregate density of FOXA1 and HNF4A motifs per site, an approach that does not test whether the relative arrangement of those motifs differs between cooperative and single-TF-enabled sites. Recent work applying deep learning to cis-regulatory sequences has shown that the spatial arrangement of transcription factor binding motifs (spacing, orientation, and relative positioning) encodes functional information beyond motif identity alone. Avsec et al. introduced the concept of “soft motif syntax” to describe distance-dependent cooperative binding rules learned by base-resolution models, including helical periodicity preferences for Nanog and directional cooperativity between pluripotency factors^7^. Liu et al. extended this framework, distinguishing hard syntax (fixed spacing below 20 bp, reflecting direct protein-protein interactions) from soft syntax (flexible spacing of 20–150 bp, consistent with nucleosome-mediated cooperativity), and classified the FOX–HNF4A composite motif in hepatocytes as soft syntax^8^. These rules were derived from the Human Development Multiomic Atlas (HDMA), a single-cell atlas of paired chromatin accessibility and gene expression spanning 817,740 cells across 12 human fetal organs, which we draw on later to test the FOXA1–HNF4A spacing grammar in primary human tissue.

The biophysical basis for soft syntax was proposed by Mirny, who showed that competition between transcription factors and a nucleosome for the same DNA region produces strongly cooperative binding without direct TF-TF interactions. This nucleosome-mediated mechanism accommodates flexible arrangements of binding sites within the nucleosomal footprint of approximately 150 bp, naturally producing the soft-syntax distance range^9^. Direct in vivo support came recently from Sönmezer et al., who used single-molecule footprinting in mouse embryonic stem cells to show that pairs of transcription factors frequently co-occupy the same DNA molecule at nucleosome-occupied cis-regulatory elements, with co-occupancy declining with inter-motif distance and independent of precise motif arrangement or physical TF-TF contact^10^.

Here, we extend Hansen’s analysis to ask whether cooperative sites carry a distinctive motif syntax, and what biochemistry underlies it. We stratify Hansen’s dual-induction CUT&Tag into four classes of co-bound sites by their dependence on single-TF expression, and identify a Cooperative class (n = 1,824) that starts in closed chromatin and opens most upon co-induction. A dual-head binding CNN trained on the four classes finds no new motif at Cooperative sites, but motif-spacing analysis reveals an enrichment of close-spaced FOXA1–HNF4A pairs (15–60 bp) specific to this class, within the soft-syntax range Avsec et al. and Liu et al. proposed for nucleosome-mediated cooperativity. We then test this nucleosomal interpretation in vitro on defined nucleosomes, using Pioneer-seq, a multiplexed nucleosome-binding assay outside the K562 cellular context. FOXA1 binds nucleosomal DNA more strongly than HNF4A per site across three reconstituted templates; and on natural nucleosomes carrying both motifs, the FOXA1 motif’s distance from the dyad predicts cobinding strength more strongly than the HNF4A motif’s. Together, the data are consistent with positioned nucleosomes acting as a scaffold for FOXA1–HNF4A cobinding, rather than only as a barrier that one factor must clear for the other.

## Results

### Four chromatin-distinct categories of co-bound FOXA1–HNF4A sites

To ask what sequence features distinguish cooperative FOXA1–HNF4A sites from those where a single factor suffices, we reanalyzed CUT&Tag data from K562 cells in which FOXA1 and HNF4A are doxycycline-induced individually or together. K562 cell line is an advantageous choice for these experiments because FOXA1 and HNF4A are not expressed in this cell lineage, making their chromatin naïve to the influence of their binding at specific sites. We took sites bound by both factors in the dual-induction condition and classified them by their dependence on single-TF expression: FOXA1-Enabled (FE, n = 1,510; bound by FOXA1 even when HNF4A is absent), HNF4A-Enabled (HE, n = 2,727; bound by HNF4A even when FOXA1 is absent), Cooperative (CB, n = 1,824; neither factor binds alone), and Redundant (n = 1,875; both factors bind independently in single-TF conditions). Overlap was called at 50% reciprocal overlap on narrowPeak intervals. Example CUT&Tag tracks from each category (Fig. 1a) show the single-condition binding logic behind the classification.

**Figure 1.**
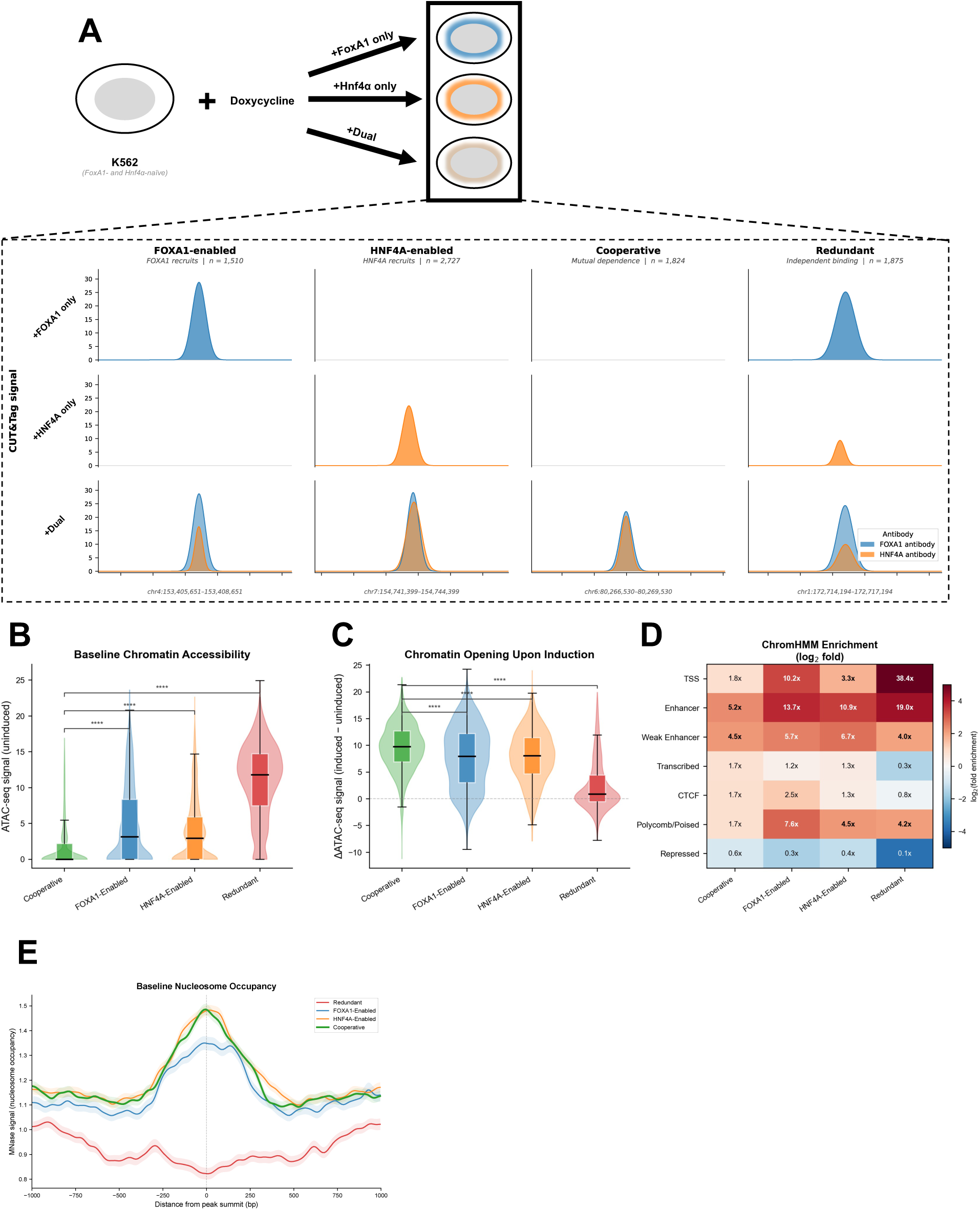
Co-bound FOXA1–HNF4A sites in K562 partition into four chromatin-distinct categories. (a) Top: schematic of the K562 doxycycline-inducible system from Hansen et al. (2022a), in which FOXA1 and HNF4A were induced individually or together in cells lacking endogenous expression of either factor. Bottom: representative CUT&Tag tracks at one peak from each category, showing FOXA1 antibody signal (blue) and HNF4A antibody signal (orange) across the three induction conditions. Co-bound sites (peaks present in both dual-induction antibody tracks; 50% reciprocal overlap on narrowPeak intervals) were classified by their dependence on single-TF expression. FOXA1-enabled (FE, n = 1,510): bound by FOXA1 in the FOXA1-only condition. HNF4A-enabled (HE, n = 2,727): bound by HNF4A in the HNF4A-only condition. Cooperative (CB, n = 1,824): bound by neither factor in either single-TF condition. Redundant (n = 1,875): bound by both factors in their respective single-TF conditions. (b) Per-peak baseline (uninduced) ATAC-seq signal by category. ATAC-seq from GSE182188, same K562 doxycycline-inducible system. (c) Change in per-peak ATAC-seq signal upon dual induction (ΔATAC = induced − uninduced). Dashed line: no change. In b and c, box plots show median (centre line), interquartile range (box), and 1.5×IQR whiskers; violins show the underlying data distribution. Brackets show two-sided Mann–Whitney U tests comparing Cooperative against each other category (****p < 0.0001). (d) Log₂ fold-enrichment of each site category over genome-wide background across seven summary chromatin states consolidated from the Broad 15-state ChromHMM K562 segmentation (wgEncodeBroadHmm). Fold enrichment = (fraction of category overlapping state) / (genomic fraction of state). Cell values are fold enrichments; colour, log₂(fold enrichment). (e) Mean MNase-seq nucleosome occupancy in a ±1 kb window centred on each peak summit, by category. MNase-seq from Mieczkowski et al. 2016 (GEO GSM2083140)^39^. Lines show category means; shaded bands show ±SEM. Sites with usable bigWig coverage (≥50% non-NaN bins): FE n = 1,420; HE n = 2,560; CB n = 1,781; RD n = 1,822. Signal binned at 10 bp and Gaussian-smoothed (σ = 20 bp).

The four categories occupy distinct chromatin contexts in K562. In uninduced cells, baseline ATAC-seq signal is near zero at cooperative sites, significantly lower than at FOXA1-Enabled or HNF4A-Enabled sites (both p < 0.0001, Mann–Whitney), while Redundant sites are constitutively open (median signal ≈ 12; Fig. 1b). Induction produces the reciprocal pattern: ΔATAC is largest at cooperative sites (median Δ ≈ 9.77), intermediate at FOXA1-Enabled and HNF4A-Enabled (median Δ ≈ 8), and minimal at Redundant sites, where baseline accessibility leaves little room to open further (Fig. 1c). ChromHMM 15-state segmentation^11,12^ of the baseline K562 epigenome separates the classes by regulatory state. Redundant sites are overwhelmingly TSS- and active-enhancer-associated (38.4× TSS enrichment, 19.0× active enhancer); cooperative sites are enhancer-biased but also enriched for polycomb-poised states; the single-TF-enabled classes fall in between (Fig. 1d). Baseline K562 MNase-seq fits this picture: the three TF-bound classes carry a positioned central nucleosome at the peak summit, while Redundant sites show a central depletion (Fig. 1e).

The same separation appears in genomic position and nearby gene expression. Cooperative sites are the most distal to transcription start sites (median 17–21 kb to nearest refGene TSS) and lie near the lowest-expressed genes in uninduced K562 (before dox; median log₂(TPM + 1) ≈ 2.5; RNA-seq from GSE182190), significantly lower than at FOXA1-Enabled sites (p < 0.0001) and indistinguishable from HNF4A-Enabled sites (Supplementary Fig. 1). Redundant sites are promoter-proximal (median ≈ 1–2 kb to nearest TSS) and lie near the highest-expressed genes (p < 0.0001 vs. Cooperative). Histone-modification metaprofiles (ENCODE K562 baseline) fit this picture. H3K4me3 and H3K27ac are sharply concentrated at Redundant sites, consistent with their promoter-proximal position. H3K4me1 is broadly enriched across all four classes, consistent with enhancer-like architecture. H3K9me3 is uniformly depleted, so none of the classes lie in constitutive heterochromatin (Supplementary Fig. 2). The four categories thus capture chromatin-distinct groups of co-bound sites in K562. Cooperative sites in particular are defined by mutual TF dependence: neither factor binds alone, and they start in closed chromatin that opens most upon co-induction.

### A dual-head binding CNN learns FOXA1 and HNF4A sequence grammar from each category

We trained a multi-task convolutional neural network with two task-specific prediction heads, one for FOXA1 and one for HNF4A, to test whether local DNA sequence is sufficient to explain binding in each of the four categories (Fig. 2a). The model takes 1,001 bp sequences centered on peak summits and outputs per-head binding probabilities. Training used 138,489 sequences: peaks from all four co-bound categories plus cis-regulatory negatives sampled from K562 DNase-accessible regions that did not overlap any FOXA1 or HNF4A peak. Training, validation, and test sets were split by chromosome to prevent leakage from genomically adjacent peaks. Held-out performance was high for both tasks. The FOXA1 head reached AUROC = 0.868 and AUPRC = 0.792; the HNF4A head reached AUROC = 0.878 and AUPRC = 0.731 (Fig. 2b).

**Figure 2.**
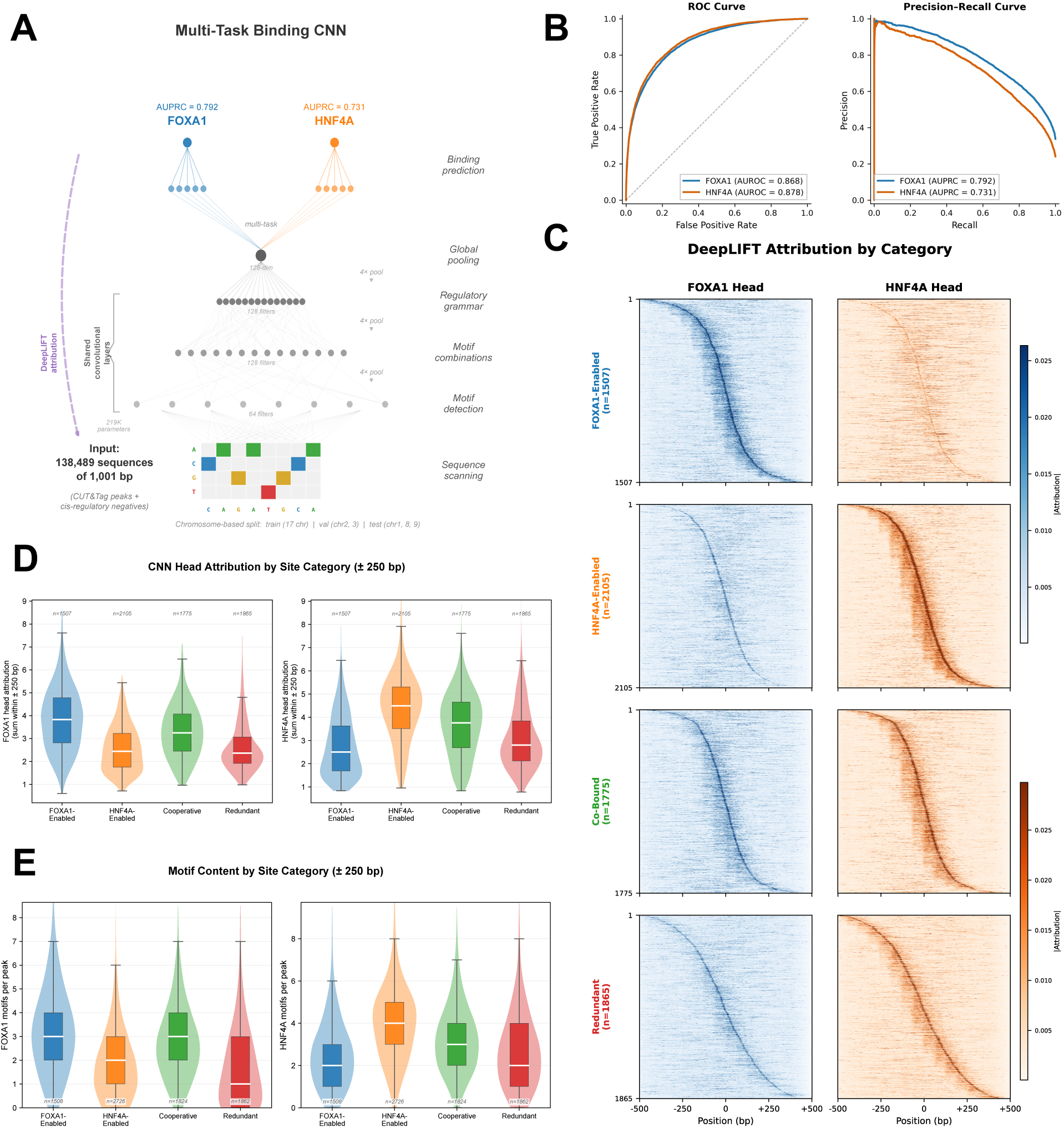
A dual-head CNN learns FOXA1 and HNF4A binding from sequence; attribution and motif content recapitulate the four categories. (a) Dual-head binding CNN architecture. One-hot encoded 1,001 bp sequences (summit ± 500 bp) pass through three convolutional blocks (64/128/128 filters; kernel sizes 19/11/7; each: Conv → BatchNorm → ReLU → MaxPool(4) → Dropout 0.25), global average pooling, and two independent task-specific MLP heads with sigmoid output. Training: 138,489 sequences (peaks from all four categories vs. cis-regulatory negatives from uninduced K562 ATAC-seq); chromosome-based splits (test: chr1, chr8, chr9; validation: chr2, chr3). (b) ROC (left) and precision-recall (right) on the held-out test set. FOXA1 head: AUROC = 0.868, AUPRC = 0.792; HNF4A head: AUROC = 0.878, AUPRC = 0.731. (c) DeepLIFT attribution heatmaps by category, after SVA filtering of the HNF4A-Enabled set (see Supplementary Fig. 4): FOXA1-Enabled (n = 1,507), HNF4A-Enabled (n = 2,105; 622 SVA-overlapping sites removed), Co-Bound (n = 1,775), Redundant (n = 1,865). Left: FOXA1 head importance (blue); right: HNF4A head importance (orange). Each row is one site; rows are sorted by position of peak attribution. Each head’s attribution is strongest at its single-TF-enabled category; both heads contribute at Co-Bound sites. (d) Total CNN head attribution within ±250 bp of the peak summit by category. FOXA1 head (left) is most active at FOXA1-Enabled sites (median 3.83 vs. 2.44 at HNF4A-Enabled); HNF4A head (right) is most active at HNF4A-Enabled sites (median 4.49 vs. 2.50 at FOXA1-Enabled). The per-site cognate-head attribution fraction (cognate-head attribution / total attribution) is higher at HNF4A-Enabled than FOXA1-Enabled sites (63.7% vs. 58.4%; two-sided Mann–Whitney p = 2.3 × 10⁻²⁴). (e) FIMO-based motif counts within ±250 bp of the peak summit (FIMO p < 10⁻³; JASPAR MA0148.1, MA0114.2). FOXA1-Enabled sites carry more FOXA1 motifs (median 3) than HNF4A motifs (median 2); HNF4A-Enabled sites show the reverse (median 4 vs. 2). The cognate-motif fraction is correspondingly higher at HNF4A-Enabled sites (71.4% vs. 57.1%; p = 7.8 × 10⁻⁸⁷). In d and e, box plots show median (centre line), interquartile range (box), and 1.5×IQR whiskers; violins show the underlying data distribution.

We then used DeepLIFT attribution^13^ ^14^ to localize which sequence positions drove each head’s prediction at each peak. Attribution heatmaps, sorted by the position of peak attribution within each category, initially showed an unexpected pattern at HNF4A-Enabled sites. Rather than a single concentrated diagonal like the one seen at FOXA1-Enabled, Cooperative, and Redundant sites, HE sites showed a bimodal “split diagonal” with two distinct attribution peaks separated by roughly 100 bp (Supplementary Fig. 4, left). We wondered whether the bimodality reflected a secondary cooperative motif grammar specific to HE sites or stereotyped sequence content producing apparent motif co-occurrence. Checking repeat-element overlap across the four categories pointed to the latter. SVA retroelements, primate-specific hominoid mobile elements^15^, whose consensus contains tandem-repeat regions with both forkhead-like and nuclear-receptor-like sub-sequences, overlapped 22.3% of HE sites (622 of 2,727) but fewer than 3% of sites in any other category (Supplementary Fig. 3). No other repeat family showed comparable class specificity. The 622 SVA-overlapping HE sites, examined alone, produced a vertical stripe barcode in attribution that matched the fixed intra-SVA positions of the composite motif features (Supplementary Fig. 4, right). Removing these 622 sites resolved the HE attribution to a single clean diagonal (Supplementary Fig. 4, middle). All subsequent attribution and motif analyses use this SVA-filtered dataset.

With SVA sites excluded, DeepLIFT attribution was consistent with each category’s binding logic. The FOXA1 head showed the strongest summit-centered attribution at FOXA1-Enabled sites; the HNF4A head was strongest at HNF4A-Enabled sites; both heads were active at Cooperative sites (Fig. 2c,d). Motif content within ±250 bp of peak summits, scored by FIMO^16^ against JASPAR 2024^17^ PWMs (MA0148.1 for FOXA1, MA0114.2 for HNF4A), tracked attribution. FOXA1 motif counts were highest at FOXA1-Enabled sites, HNF4A counts were highest at HNF4A-Enabled sites, and Cooperative sites carried both (Fig. 2e). Attribution and motif content agreed, and neither identified a qualitatively new sequence feature at Cooperative sites beyond the presence of both canonical motifs.

To test more rigorously whether Cooperative sites contain any de novo composite or otherwise distinctive pattern that attribution could be missing, we ran TF-MoDISco^18^ on the SVA-filtered per-head attributions, splitting by category for eight independent runs (four categories × two heads). Every run recovered the canonical motif for its head, forkhead for the FOXA1 head and nuclear-receptor direct-repeat (DR1) for the HNF4A head, in every category (Supplementary Fig. 5). No novel cooperative-specific motif emerged. If anything distinguishes cooperative sites from single-TF-enabled sites at the sequence level, it is not a new motif but the relative arrangement of the canonical two motifs.

### Cooperative sites carry close-spaced FOXA1–HNF4A motif pairs

To ask whether the arrangement of the canonical motifs differs between the four categories, we scanned each peak (summit ± 500 bp) with FIMO (p < 10⁻³) using JASPAR 2024 PWMs MA0148.1 (FOXA1) and MA0114.2 (HNF4A), and computed per-peak FOXA1–HNF4A center-to-center spacing using a best-score pairing strategy (lowest-pvalue FOXA1 motif paired with lowest-pvalue HNF4A motif, max distance 500 bp). Cooperative sites had the shortest median spacing (146 bp; n = 1,599 peaks with both motifs), relative to FOXA1-Enabled (180 bp; n = 1,277), HNF4A-Enabled (182 bp; n = 1,827; SVA-filtered), and Redundant (197 bp; n = 1,295) (Fig. 3a).

**Figure 3.**
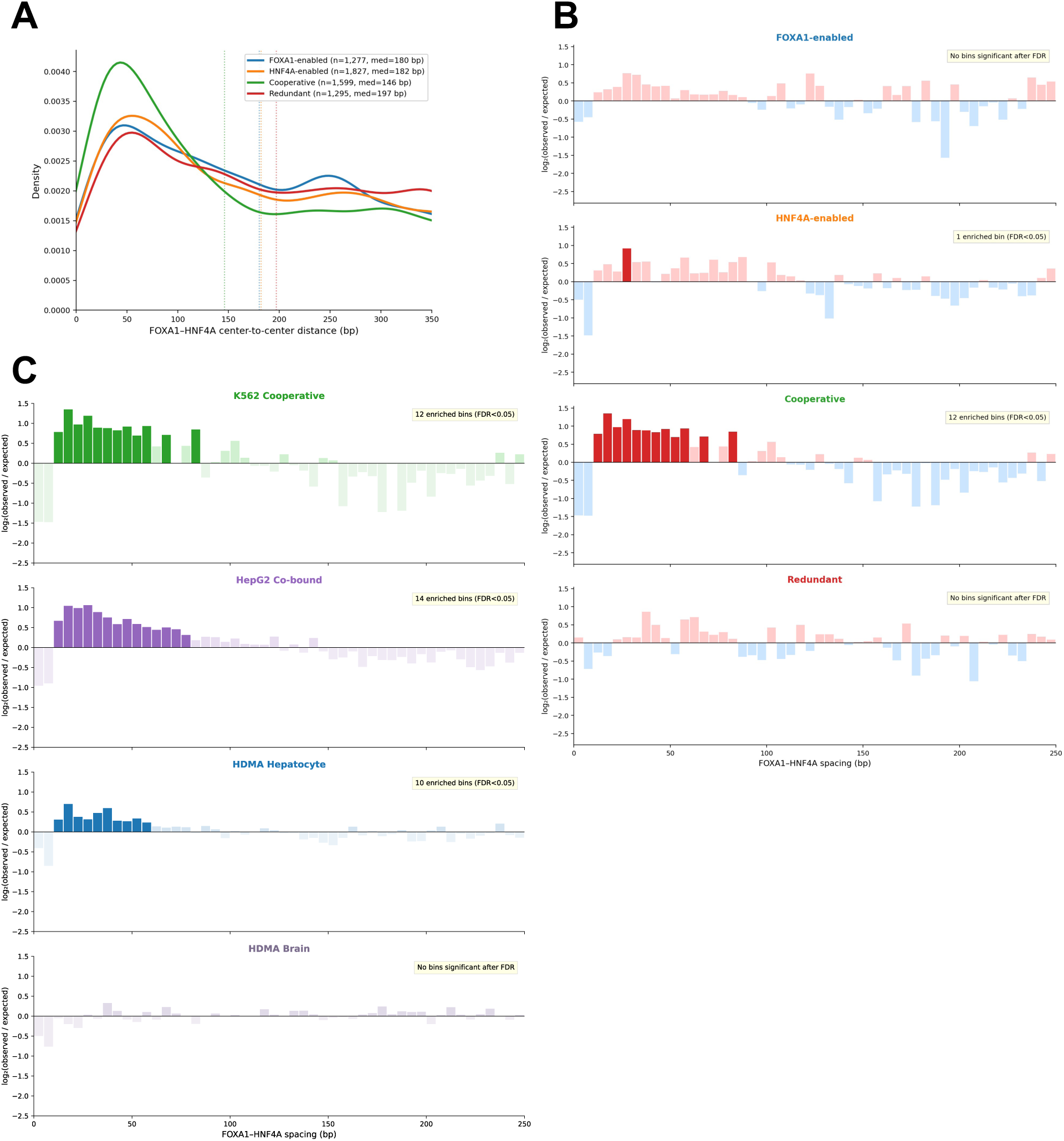
Cooperative sites carry close-spaced FOXA1–HNF4A motif pairs absent from sites where one factor suffices. Spacing: centre-to-centre between lowest-p FOXA1 (MA0148.1) and HNF4A (MA0114.2) motifs per peak, summit ± 500 bp, max 500 bp inter-motif (best-score pairing; FIMO p < 10⁻³). (a) Kernel density of per-peak motif spacing by category; lines mark medians. Cooperative is shortest (146 bp) vs FOXA1-enabled (180), HNF4A-enabled (182; SVA-filtered), Redundant (197); n = peaks with both motifs per category. (b) Per-bin log₂(observed/expected) in 5 bp bins against a 1,000-permutation per-peak null (motif positions shuffled within the 1,001 bp window). Dark red, FDR-enriched (BH q < 0.05); dark blue, depleted; pale, n.s. Cooperative shows 12 enriched bins at 15–60 bp; FOXA1-enabled and Redundant show 0; HNF4A-enabled shows 1 (Suppl. Fig. 6 for unfiltered). (c) Same pipeline at endogenously co-bound sites. Top: K562 Cooperative, replotted from (b). Second: HepG2 (FOXA1–HNF4A ChIP-seq, GSE104247; 9,373 motif pairs), 14 enriched bins. Third: HDMA fetal hepatocyte caCREs (Liu et al. 2026, Nature; 44,165 peaks in clusters LI_1/3/4/6 from 29,926 cells, PCW15–22; 24,327 motif pairs), 10 enriched bins concentrated at 15–60 bp. Bottom: HDMA fetal brain caCREs (BR_0–BR_17; 74,035 peaks, 14,015 motif pairs), 0 enriched bins.

To ask whether specific inter-motif distances are enriched beyond what random motif co-occurrence within a peak would produce, we computed log₂(observed / expected) per 5-bp spacing bin against a permutation null (1,000 shuffles of motif positions within the 1,001-bp FIMO search window, best-score pairing applied identically to observed and null). At Cooperative sites, ten contiguous 5-bp bins spanning 15–60 bp were significantly enriched after Benjamini–Hochberg FDR correction (q < 0.05) (Fig. 3b). At HNF4A-Enabled sites (SVA-filtered), a single isolated bin at 25–30 bp reached significance (q = 4.6 × 10⁻³), but no other bins in the 15–60 bp range did. No bin in this range was enriched at FOXA1-Enabled or Redundant sites. The contiguous close-spacing signal is most specific to Cooperative sites.

We tested whether this signal could be explained by residual repeat-element content not captured by the SVA filter. Ranking repeat families by their CB category-specific enrichment z-score (Supplementary Fig. 3), we iteratively removed peaks overlapping the three most CB-enriched families (Alu, z = +1.39, 19.5% of CB peaks; ERVL, z = +1.30, 5.4%; ERVK, z = +1.22, 1.9%) and re-ran enrichment on each reduced set. Six, nine, and ten bins respectively remained FDR-enriched in the 15–60 bp range after removal of each family (Supplementary Fig. 7), and the peak of the enrichment remained in that range in every case. The reduction in bin count tracks the loss of statistical power from smaller samples rather than a shift in the spacing signal.

Before SVA filtering, HNF4A-Enabled sites carried an apparent enrichment of 2 bins at ∼95–105 bp (Supplementary Fig. 6). These bins overlap the characteristic inter-motif distance within the SVA composite element, and removing the 622 SVA-overlapping HE peaks eliminated this enrichment (leaving 1 bin significant). The HE spacing profile shown in Fig. 3b is the SVA-filtered version. That SVA contamination was a class-level feature of HE rather than a peculiarity of the 622 SVA peaks alone is consistent with CentriMo^19^ analysis of the full HE set, which showed the canonical FOXA1 motif to be only weakly centrally enriched at HE peak summits, substantially less so than the related forkhead FOXK1 motif (Supplementary Fig. 8). The forkhead-like signal at HE sites was therefore not FOXA1 binding per se. The SVA-overlapping HE sites are themselves legitimate HNF4A-enabled co-binding events; they are excluded from spacing analysis because their fixed composite architecture determines the inter-motif distance by repeat sequence, not by a functional spacing preference.

To test whether the close-spacing grammar generalizes beyond the K562 dox-induction system, we re-applied the per-bin spacing analysis across settings of progressively greater biological realism (Fig. 3c). First, FOXA1–HNF4A co-bound peaks in the HepG2 hepatoma cell line, where both factors are endogenously expressed (ChIP-seq from GSE104247; 9,373 motif-containing peaks), showed 14 FDR-enriched bins at 15–60 bp, overlapping the K562 Cooperative enrichment range. We then moved from cancer lines to primary tissue using HDMA, a single-cell multiomic atlas of 12 human fetal organs^8^. Fetal liver at post-conception weeks 15–22 is the native developmental context for FOXA1 and HNF4A, which jointly specify hepatocyte identity, and the atlas’s single-cell resolution lets us test genuine hepatocyte regulatory elements rather than a bulk-tissue average. We applied the same pipeline to chromatin-accessible candidate cis-regulatory elements (caCREs) from the atlas’s hepatocyte clusters LI_1, LI_3, LI_4, and LI_6 (29,926 cells; 44,165 caCREs; 24,327 motif-containing peaks; Supplementary Fig. 9). Hepatocyte caCREs reproduced the close-spacing enrichment with 10 FDR-enriched bins concentrated at 15–60 bp, despite a smaller per-peak motif yield from the tissue-derived peak set. Close-spaced peaks in the same hepatocyte set were also more accessible than far-spaced peaks (1.20-fold; one-sided Mann–Whitney p = 2.3 x 10^-11; Supplementary Fig. 10), consistent with cooperative occupancy at the close-spaced subset. As a matched negative control from the same atlas, we ran the identical pipeline on fetal brain caCREs (clusters BR_0 through BR_17; 74,035 caCREs; 14,015 motif-containing peaks), where neither FOXA1 nor HNF4A is appreciably expressed at this developmental stage. Brain caCREs showed zero FDR-enriched bins anywhere in the 0–250 bp range.

The Cooperative close-spacing grammar is therefore specific to this category, robust to removal of CB-enriched repeat families, reproducible in both an unrelated cancer line (HepG2) and primary fetal hepatocyte tissue, and absent from a FOXA1/HNF4A-negative fetal brain control. The pattern is not a generic property of motif-rich active chromatin but a tissue-context-specific signature of FOXA1–HNF4A co-binding. Its 15–60 bp range is within the soft-syntax window (15–150 bp) that Avsec et al. 2021 and Liu et al. 2026 proposed as consistent with nucleosome-mediated cooperativity^9^.

### Pioneer-seq shows FOXA1 binds nucleosomes more strongly than HNF4A across templates

To test whether the single-TF binding asymmetry Hansen et al. observed in cells reflects an intrinsic difference in nucleosome-binding affinity, we used Pioneer-seq, a multiplexed in vitro nucleosome-binding assay^20,21^. The library places a single FOXA1 or HNF4A binding site at each of 182 positions (−85 to +96 bp relative to the dyad) on three reconstituted nucleosomal templates (Widom 601, 5S, MMTV)^22^ (Fig. 4a). A nonspecific sequence (a partial ETS motif, JASPAR MA0098.3) was placed at matched positions on the same templates as a paired control.

**Figure 4.**
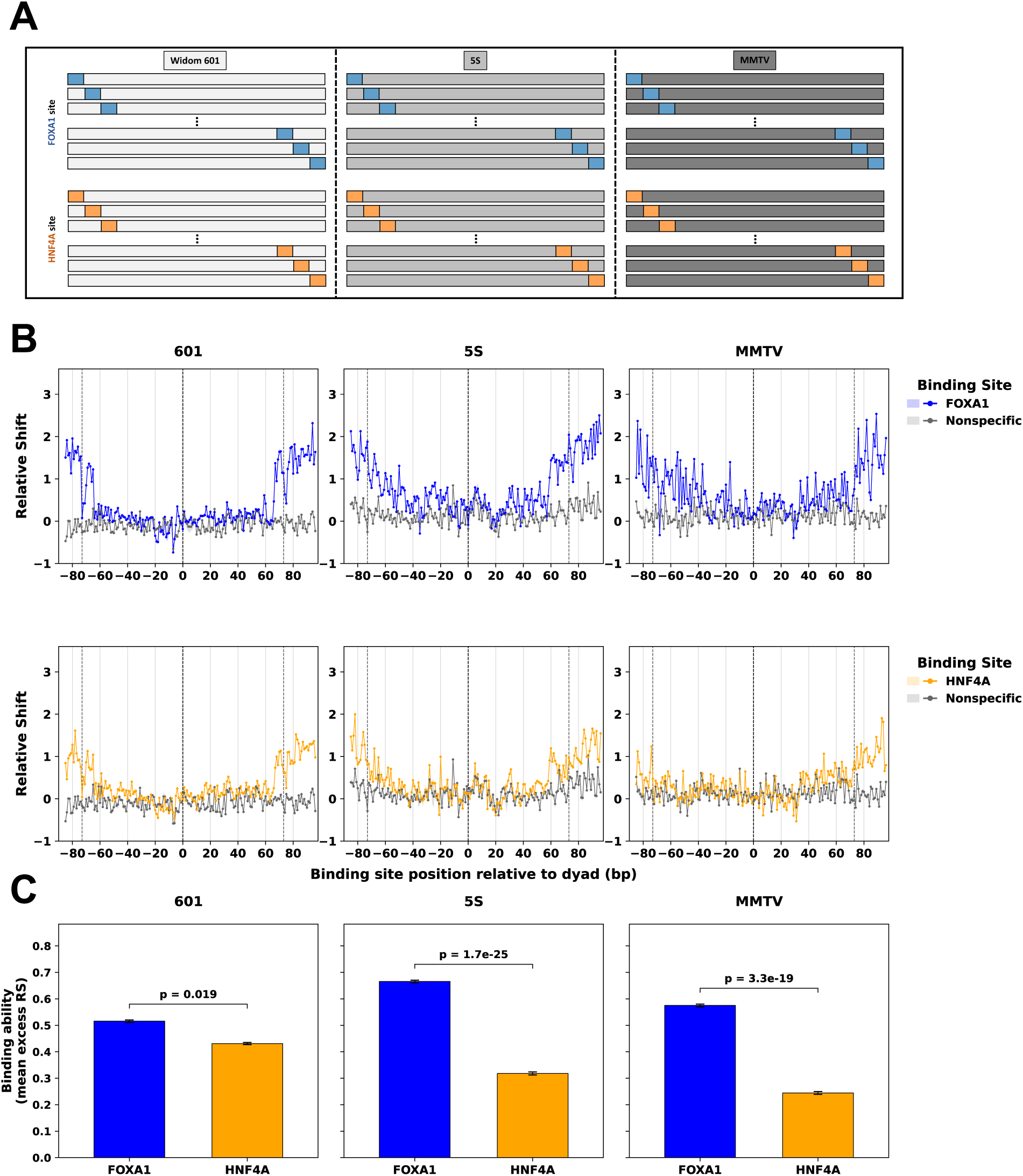
FOXA1 binds nucleosomes more strongly than HNF4A across three nucleosomal templates. (a) Single-site Pioneer-seq library design. A single FOXA1 binding site (blue; TGTTTACTTTG, JASPAR MA0148.1) or a single HNF4A binding site (orange; GAGTCCAAAGTCCAG, JASPAR MA0114.2) was placed at each of 182 centre positions (−85 to +96 bp relative to the dyad) on three reconstituted nucleosomal templates: Widom-601, 5S rDNA, and mouse mammary tumor virus (MMTV)-A. A paired nonspecific control sequence (a partial ETS motif; ACCGGAAGTG, JASPAR MA0098.3) was placed at matched positions on the same templates. Each row of the schematic represents one library member. (b) Relative shift (RS) as a function of binding-site centre position relative to the nucleosome dyad. Top row: FOXA1 (blue) and the paired nonspecific control (grey). Bottom row: HNF4A (orange) and the paired nonspecific control. Points show the mean of n = 3 biological replicates; vertical error bars, SEM. Vertical dashed lines mark the dyad (position 0) and the canonical nucleosome boundaries (±73 bp). RS is defined as −log₂((T / T_NS) / (N / N_NS)), where T and T_NS are read counts of the test and paired nonspecific-control nucleosomes in the unshifted band of the TF-treated lane, and N and N_NS are the corresponding counts in the no-TF (null) lane (Methods). (c) Binding ability per template, defined as the mean excess RS over the nonspecific control, ⟨RS_TF − RS_NS⟩, averaged across all 182 positions. Bars show the mean; error bars, SEM propagated from per-position SEMs. p-values, one-sided paired Wilcoxon signed-rank test (alternative: FOXA1 > HNF4A; 182 paired positions per template); the directional hypothesis was prespecified from cellular observations (Hansen et al., 2022a) of FOXA1’s lower per-motif binding requirement.

Relative-shift (RS) profiles, computed as the log-ratio depletion of each test sequence from the unshifted band of the TF-bound lane versus the matched nonspecific control (see Methods), show that both FOXA1 and HNF4A bind nucleosomal DNA above nonspecific at multiple positions across all three templates (Fig. 4b). For both TFs, RS rises near the nucleosome ends and at linker-like positions where DNA is more accessible. Across templates, FOXA1 produced higher RS than HNF4A at most positions. Quantifying per-TF binding ability as the mean excess RS over the nonspecific control averaged across all 182 positions, ⟨RS_TF − RS_NS⟩, FOXA1 exceeded HNF4A on all three templates (Fig. 4c): 601 (0.52 vs 0.43; paired Wilcoxon p = 0.019), 5S (0.67 vs 0.32; p = 1.7 × 10⁻²⁵), and MMTV (0.58 vs 0.24; p = 3.3 × 10⁻¹⁹). The effect was largest on the 5S and MMTV templates and smallest on the engineered Widom 601. Raw EMSA titration gels for all three biological replicates underlying the quantification are shown in Supplementary Fig. 8. The single-TF binding difference Hansen observed in cells is therefore mirrored in vitro: across three independent nucleosomal templates and 182 positions per template, FOXA1 engages nucleosomal DNA more strongly than HNF4A at most positions tested.

### Cobinding on nucleosomes synergizes and tracks FOXA1 position

To test whether the per-nucleosome affinity asymmetry from Figure 4 affects cobinding, we built a second Pioneer-seq library in which a FOXA1 and an HNF4A binding site were placed adjacently with a fixed 5-bp gap between the sites, at each of 77 dyad-distance positions on the Widom 601, 5S, and MMTV templates (Fig. 5a). The 5-bp gap matches the most enriched inter-motif spacing at K562 Cooperative sites (Fig. 3b) and is independently supported by Spaced Motif Analysis (SpaMo, MEME Suite)^23^. Running SpaMo on Cooperative peak sequences with FOXA1 (MA0148.1) as the primary motif, HNF4A was the top-scoring secondary motif at a best gap of 5 bp upstream in primary-palindromic orientation (secondary PWM MA1494.2, an alternative JASPAR HNF4A profile to MA0114.2 built from an independent dataset; E = 4.24 × 10⁻⁴; Supplementary Fig. 11).

**Figure 5.**
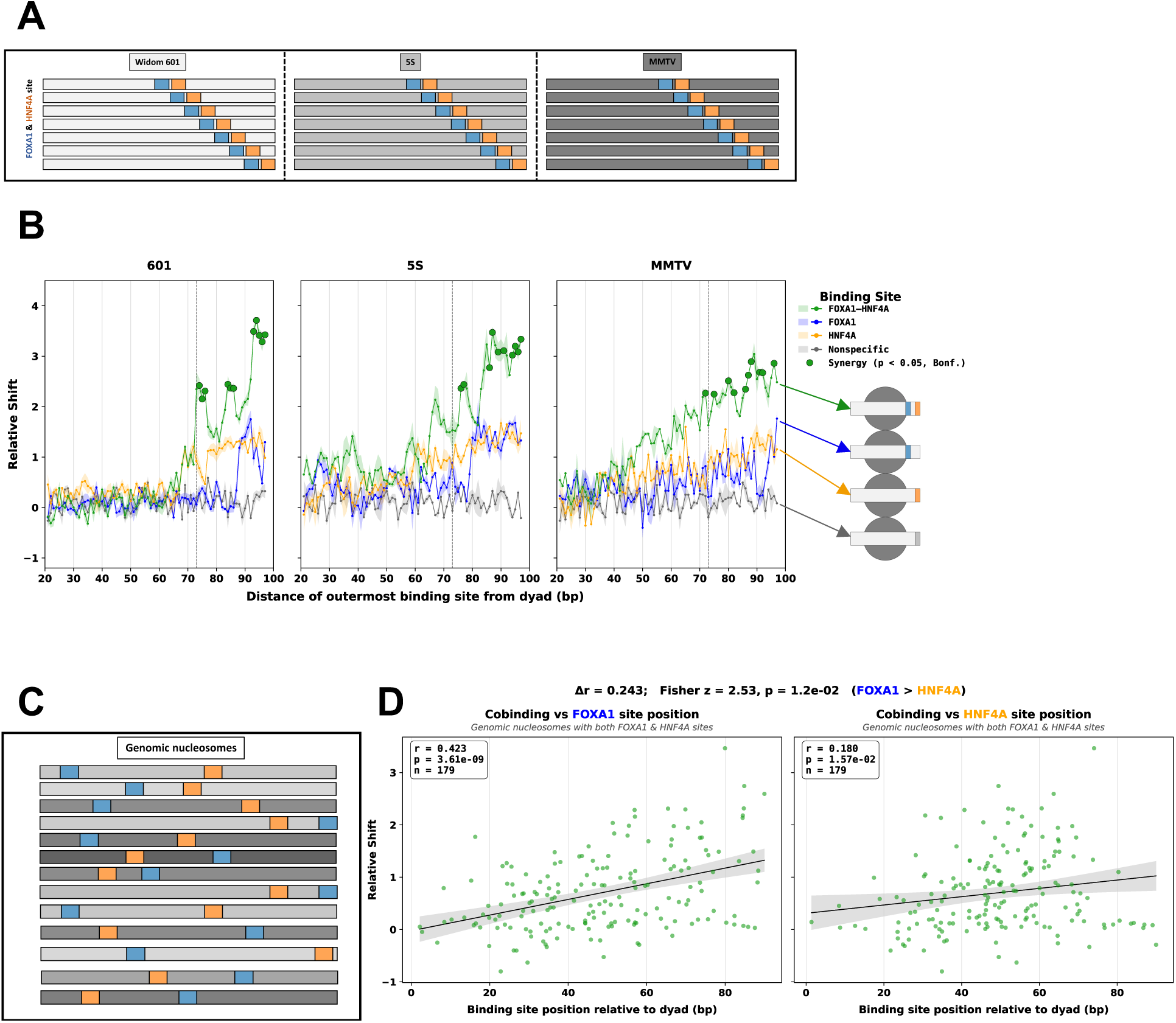
FOXA1–HNF4A cobinding synergises on defined nucleosomes and, on natural nucleosomes, tracks the FOXA1 motif’s dyad position more than the HNF4A motif’s. (a) Cobinding Pioneer-seq library design. On each of three nucleosomal templates (Widom-601, 5S rDNA, mouse mammary tumor virus (MMTV)-A; light-to-dark grey shading), a FOXA1 site (blue) and an HNF4A site (orange) were placed adjacently with a fixed 5 bp gap between the two sites, at each of 77 outermost-site positions (bp 21–97 from the dyad). Each row in the schematic represents one library member. (b) Pioneer-seq relative shift (RS) as a function of the outermost site’s distance from the dyad. Green: FOXA1–HNF4A composite (TGTTTACTTTG–N₅–GAGTCCAAAGTCCAG; JASPAR MA0148.1 + MA0114.2). Blue: FOXA1 alone (MA0148.1). Orange: HNF4A alone (MA0114.2). Grey: paired nonspecific control (ACCGGAAGTG; JASPAR MA0098.3). Per-position values are the mean of n = 3 biological replicates with SEM error bars. Vertical dashed line marks the canonical nucleosome edge (bp 73). Large green dots mark positions where the cobinding signal exceeds the sum of single-TF signals on the linear scale (2^FH > 2^F + 2^H; paired z-test with delta-method error propagation; Bonferroni-corrected across the 77 positions per template, α = 0.05). Cartoons at right depict the four binding conditions, colour-matched to the trace lines. (c) Genomic-nucleosome library. Each row represents one of n = 179 nucleosomes selected from K562 Cooperative-category peaks containing exactly one FOXA1 motif (blue) and exactly one HNF4A motif (orange) (FIMO p < 10⁻³, JASPAR MA0148.4 for FOXA1 and MA0114.4 for HNF4A; nucleosome dyads inferred by DANPOS from K562 MNase-seq (Mieczkowski et al. 2016) at occupancy score ≥ 0.7; Methods). (d) Cobinding RS on the genomic-nucleosome library versus the FOXA1 motif’s distance from the inferred dyad (left) and the HNF4A motif’s distance from the inferred dyad (right). Points, individual nucleosomes; line, ordinary least squares fit; grey band, 95% CI. In-panel: Pearson r, two-sided p, and n. Banner: Δr = r_FOXA1 − r_HNF4A and two-sided Fisher z-test comparing the two Pearson correlations.

Relative-shift profiles with both TFs co-titrated (green) substantially exceed either single-TF profile (FOXA1, blue; HNF4A, orange) at a cluster of positions toward the nucleosome ends on each template (Fig. 5b). Per-position synergy was tested as 2^FH > 2^F + 2^H by paired z-test with delta-method error propagation on the linear scale (Bonferroni-corrected across 77 positions, α = 0.05); significant synergy concentrated at bp 72–97 on each template.

To test whether this synergy on engineered templates extends to natural sequences with their inherent positional variability, we assayed a second library of n = 179 genomic nucleosomes that natively contain both a FOXA1 and an HNF4A motif (Fig. 5c). Cobinding RS on this library correlated with the position of the FOXA1 motif relative to the inferred nucleosome dyad (Pearson r = 0.423, p = 3.6 × 10⁻⁹) more strongly than with the HNF4A motif’s dyad distance (r = 0.180, p = 1.6 × 10⁻²) (Fig. 5d). The two correlations differ significantly (Δr = 0.243; Fisher z = 2.53, two-sided p = 1.2 × 10⁻²).

The asymmetry that predicts cobinding strength on these natural nucleosomes is therefore in the FOXA1 motif’s position, not the HNF4A motif’s. This does not mean the two motifs need to occupy the same nucleosomal face: many of the highest-cobinding nucleosomes carry FOXA1 and HNF4A motifs on opposite sides of the dyad, suggesting the dominant constraint is whether the pair sits on a positioned nucleosome at all, not whether the two motifs share a face.

## Discussion

The classical pioneer factor hypothesis, formulated from Cirillo et al.’s reconstitutions of FoxA on the albumin enhancer^1,2^, held that pioneer factors clear nucleosomal barriers so non-pioneer factors can subsequently bind exposed DNA. Hansen et al. challenged this by showing that both FOXA1 and HNF4A can independently bind and open inaccessible chromatin in K562 cells, and recast pioneer activity as a quantitative property of TF–DNA affinity and motif content^5,6^. Our analysis sits between these views. At Cooperative sites, where both factors are required for binding, FOXA1 and HNF4A motifs are enriched at 15–60 bp inter-motif spacing, a syntax absent from single-TF-enabled sites and within the soft-syntax range previously proposed for TF cooperativity^7,8^. The biophysical mechanism is the one proposed by Mirny ^9^, where two transcription factors competing with the same nucleosome bind cooperatively without direct TF–TF contact, because transient unwrapping that exposes one motif also exposes its neighbor, so each factor lowers the effective barrier for its partner. The Cooperative class shows this regime’s genomic fingerprint: closed chromatin at baseline (Fig. 1b,c), opened only when both factors are present, and marked by a close-spaced FOXA1–HNF4A motif pair within a single nucleosomal footprint (Fig. 3b, Fig. 5b). The same fingerprint appears in vivo: single-molecule footprinting in mouse embryonic stem cells finds pairwise TF co-occupancy concentrated at nucleosome-occupied regions, declining with inter-motif distance and indifferent to precise motif arrangement^10^. Together with our Pioneer-seq results on defined nucleosomes (Fig. 4, Fig. 5), the data position cooperative FOXA1–HNF4A binding as a structural phenomenon, with a positioned nucleosome acting as a shared scaffold for two factors that engage it competitively rather than sequentially.

We confirm Hansen’s observation that the FOXA1–HNF4A system contains a Cooperative class of sites that requires both factors, and we further specify that this requirement has a structural sequence basis: at Cooperative sites the two TFs’ canonical motifs are enriched at 15–60 bp inter-motif spacing consistent with a soft-syntax. In this model, transient unwrapping that exposes one motif can also expose its neighbor, so each factor lowers the effective barrier for its partner, and the pair binds cooperatively without direct TF–TF contact. Our Cooperative class is the genomic signature of exactly that model, closed at baseline (Fig. 1b,c), opened only when both factors are present, and marked by a FOXA1–HNF4A motif pair within a single nucleosomal footprint (Fig. 3e, Fig. 5b). Sönmezer et al. provided direct in vivo evidence for this mechanism in mouse embryonic stem cells, showing that high pairwise TF co-occupancy occurs preferentially at nucleosome-occupied regions, declines with inter-motif distance, and does not require strict motif organization, paralleling the soft, distance-bounded grammar we observe at FOXA1–HNF4A cooperative sites. Together with our in vitro Pioneer-seq results on defined nucleosomes (Fig. 4, Fig. 5), the picture is that cooperativity at FOXA1–HNF4A sites reflects neither a categorical TF identity nor only TF–DNA affinity, but a structural arrangement of motifs on positioned nucleosomes where both factors competitively engage the same nucleosomal footprint.

The two motif-position predictors of cobinding strength on natural nucleosomes are not symmetric. Cobinding relative shift correlates with the FOXA1 motif’s dyad distance more strongly than with the HNF4A motif’s (r = 0.42 vs r = 0.18; Δr = 0.24, Fisher z p = 1.2 × 10⁻²), and FOXA1 also binds nucleosomal DNA more strongly than HNF4A at most positions across three reconstituted templates. The simplest interpretation is that FOXA1’s per-nucleosome engagement provides the limiting access, with HNF4A free to bind elsewhere on the same nucleosome. Consistent with this, many of the highest-cobinding genomic nucleosomes carry FOXA1 and HNF4A motifs on opposite sides of the dyad rather than co-positioned on the same nucleosomal face. The dominant constraint on cobinding therefore appears to be whether the pair sits on a positioned nucleosome at all, typically because FOXA1 can engage that nucleosome, rather than precise co-positioning of the two motifs. We hesitate to call FOXA1 the “lead partner” in a strict sense; the asymmetry we observe is one of two predictors, not a deterministic causal claim.

Two recent findings bear on this picture. Hansen and Cohen used doxycycline titration in the same K562 induction lines to compute a pioneer-activity index, defined as the ratio of average dox₅₀ at accessible versus inaccessible binding sites, and reported that HNF4A has a higher index (0.680) than FOXA1 (0.391)^6^. Our Pioneer-seq result that FOXA1 binds defined nucleosomes more strongly than HNF4A appears at first to disagree, but the two metrics measure different observables. Hansen and Cohen’s index integrates genome-wide motif availability, cellular cofactor recruitment, and each TF’s relative affinity for the in-cell distinction between accessible and inaccessible chromatin; Pioneer-seq measures per-nucleosome biochemistry on defined templates without those degrees of freedom. Both findings hold: per-nucleosome affinity favors FOXA1; in-cell pioneer activity by titration favors HNF4A.

Liu et al. (2026) used ChromBPNet trained on 817,000 fetal scATAC profiles to predict a lexicon of cooperative motif syntaxes, and assigned the FOX–HNF4A composite to soft syntax based on contribution-score marginalization^8^. Our results provide direct experimental measurement of that prediction at the resolution of the cooperative subset, which observational fetal data cannot isolate without single-TF perturbation. The two studies are complementary.

Another consideration for interpreting these sites concerns transposable elements. Two patterns of TE overlap appear in our data. Primate-specific SVA retroelements overlap 22.3% of Hansen et al.’s “HNF4A-enabled” category (Supplementary Fig. 3). Their consensus carries both forkhead-like and nuclear-receptor-like sub-sequences in tandem, and attribution at these sites traces fixed intra-SVA motif positions rather than a functional TF–TF arrangement. Alu, ERVL, and ERVK elements are modestly enriched at Cooperative sites (19.5%, 5.4%, and 1.9% of CB peaks respectively), consistent with the well-documented role of transposable elements as a major source of novel transcription factor binding sites in mammalian genomes and of primate-specific regulatory innovation across pluripotency, placental, and immune networks^24–28^. We interpret the TE-enriched fraction of co-bound peaks as an evolutionary byproduct of retrotransposon expansion rather than as cis-grammatical cooperativity in the sense of this study: repeat insertion generates motif co-occurrence at spacings determined by the element’s internal architecture, not by selection for cooperative TF engagement. Some of these events may subsequently have been co-opted into regulatory function, the canonical mechanism of TE-derived enhancer evolution, but distinguishing co-opted from incidental cases requires targeted functional testing beyond this study. Critically, the 15–60 bp cooperative-site spacing signal survives iterative removal of the three most-enriched repeat families, indicating that two signatures coexist in the genome and must be separated to recover the cis-grammar that specifies cooperative binding independent of retrotransposon content.

Several limitations bound these claims. The K562 CUT&Tag and ATAC-seq data we reanalyse are from cells in which FOXA1 and HNF4A are dox-induced at approximately 1,000-fold above endogenous levels^5^. The close-spacing grammar reproduces at HepG2 and in primary fetal hepatocyte caCREs at native expression (Fig. 3c), so it is not an induction artifact; but the four-class chromatin partition itself (Fig. 1) requires the K562 dual-induction system and cannot be recovered in cells where both factors are constitutively expressed. Pioneer-seq assays reconstituted nucleosomes on synthetic templates and lacks the native chromatin context that shapes binding in cells: neighboring nucleosomes, histone modifications, cofactor occupancy, and three-dimensional folding. Our cobinding analysis on genomic nucleosomes (Fig. 5d) restores native sequence variability but is still in vitro. We do not directly test whether FOXA1 must arrive before HNF4A at any cooperative site; the architecture we describe is consistent with both sequential and simultaneous engagement of a pre-positioned nucleosome. Single-molecule live-cell imaging of labeled FOXA1 and HNF4A would address this temporal question that our equilibrium assays cannot.

Three observables measure different quantities: per-motif efficiency (Hansen, 2022a)^5^, per-nucleosome affinity (this work), and in-cell pioneer activity by titration (Hansen and Cohen, 2022b)^6^. Treating them as competing answers to “which TF is the better pioneer” conflates distinct biochemistry. The classical pioneer factor model framed by Cirillo et al. (2002) holds at single-TF-enabled sites: one factor engages the nucleosome and lowers its barrier to others. At cooperative FOXA1–HNF4A sites, the data are better explained by the nucleosome as a scaffold than as a barrier one factor must clear for the other.

## Methods

### Cell lines and public data sources

CUT&Tag data for FOXA1, HNF4A, and FOXA1+HNF4A dox-induced K562 lines (0.5 µg/ml dox, 24 h induction) were obtained from GEO accession GSE182189^5^. ATAC-seq from the same induction conditions was obtained from GSE182188; RNA-seq from GSE182190. Doxycycline-titration CUT&Tag from the same lines^6^ was obtained from GSE204726 but was not used for this study’s analyses. HepG2 FOXA1 and HNF4A ChIP-seq for cross-cell-type validation (Fig. 3c) was obtained from GSE104247^29^. ChromHMM 15-state K562 segmentation, K562 DNase-seq, and ENCODE K562 baseline histone modification ChIP-seq tracks (H3K4me1, H3K4me3, H3K27ac, H3K27me3, H3K9me3) were obtained from the ENCODE portal. RepeatMasker hg19 annotations were obtained from UCSC. SVA element annotations were extracted by relabelling RepeatMasker repFamily=“Other” entries. All genomic coordinates use hg19 (GRCh37).

### Peak processing and classification

CUT&Tag narrowPeak files for each replicate were intersected at 50% reciprocal overlap to produce consensus peak sets per condition (FOXA1-alone FOXA1-antibody, HNF4A-alone HNF4A-antibody, dual-induction FOXA1-antibody, dual-induction HNF4A-antibody). Peaks falling outside the 24 standard chromosomes (chr1–22, chrX, chrY) were excluded. Co-bound sites were defined as peaks present in both dual-induction conditions, summit-centred to 1,001 bp windows. Each co-bound site was then classified by single-TF dependence using 50% reciprocal overlap with single-induction peaks: FOXA1-Enabled = also bound by FOXA1 in the FOXA1-alone condition (n = 1,510); HNF4A-Enabled = also bound by HNF4A in the HNF4A-alone condition (n = 2,727); Cooperative = bound by neither factor in either single-TF condition (n = 1,824); Redundant = bound by both factors in their respective single-TF conditions (n = 1,875). This peak-resolution classification differs from the gene-centric scheme of Hansen et al. (2022a) but uses the same primary CUT&Tag data. Cis-regulatory negatives for CNN training were sampled from ENCODE K562 DNase peaks that did not overlap any consensus FOXA1 or HNF4A peak in any condition.

### Chromatin accessibility and histone modification analysis

ATAC-seq alignment was as in Hansen et al. (2022a). Per-peak baseline ATAC and ΔATAC (induced − uninduced) signal was computed as RPKM-normalised read coverage in a ±150 bp window around the consensus peak summit. ChromHMM enrichment per category was computed as the log₂ ratio of observed-to-expected base-pair overlap with each of the 15 K562 chromatin states. Histone-mark and DNase metaprofiles (Supplementary Fig. 2) were generated with deepTools^30^ (computeMatrix reference-point --referencePoint center -a 2000 -b 2000 -bs 50; plotProfile).

### Binding CNN

A multi-task convolutional neural network was implemented in PyTorch^31^. The architecture has three shared 1D convolutional blocks (kernel sizes 19/11/7; 64/128/128 filters; each block: Conv → BatchNorm → ReLU → MaxPool(4) → Dropout(0.25)), followed by global average pooling (output dimension 128) and two parallel task-specific heads. Each head is a two-layer MLP (Linear 128→32 → ReLU → Dropout(0.25) → Linear 32→1) with sigmoid output. Total trainable parameters: ∼219,000.

Inputs are 1,001 bp DNA sequences one-hot encoded as (4, 1001) tensors. Training used the union of all four co-binding categories and cis-regulatory negatives (138,489 sequences total after deduplication and SVA filtering), with negatives capped at 2× the positive count. Train/validation/test splits were chromosome-based (test = chr1, chr8, chr9; validation = chr2, chr3; train = remaining standard chromosomes), following the BPNet convention^7^. Loss was BCE with class-balanced positive weights. Optimisation used Adam (lr = 10⁻³, weight decay = 10⁻⁴) with ReduceLROnPlateau (factor 0.5, patience 5) on validation loss, early stopping with patience 10, and a maximum of 60 epochs at batch size 128. SVA-overlapping peaks were excluded from training (initial training without exclusion produced the split-diagonal attribution artifact described in Supplementary Fig. 4).

### DeepLIFT attribution and TF-MoDISco

Per-position attribution scores were computed with DeepLIFT^13^ (Captum implementation; reference: shuffled-input dinucleotide-frequency-matched). Attributions were sorted by the position of peak attribution within each binding-site category to generate the heatmaps in Fig. 2c. TF-MoDISco^18^ (modisco-lite implementation) was run separately for each (category × head) combination with default parameters apart from max_seqlets_per_metacluster = 5,000 and n_leiden_runs = 2. The top-ranked positive contribution-weight-matrix logo from each run is shown in Supplementary Fig. 5.

The full training, inference, and attribution pipeline is available at: https://github.com/pwilson97/FOXA1-hnf4a-binding-cnn

### FIMO motif scanning and spacing analysis

Sequences (summit ± 500 bp; 1,001 bp total) were scanned with FIMO (MEME Suite v5.5.7^32^; --parse-genomic-coord --thresh 1e-3) using JASPAR 2024 PWMs MA0148.1 (FOXA1) and MA0114.2 (HNF4A). Per-peak spacing was computed as the centre-to-centre distance between the lowest-pvalue FOXA1 motif and the lowest-pvalue HNF4A motif (best-score pairing strategy), with maximum inter-motif distance 500 bp. Permutation null distributions were generated by shuffling motif positions within each peak’s 1,001 bp FIMO search window (1,000 permutations, seed 42) and computing best-score pair distances per shuffle. Per-bin enrichment was computed as log₂(observed / expected) in 5 bp bins, with bin-wise p-values from the permutation distribution and Benjamini–Hochberg FDR correction across bins (q < 0.05).

### HDMA cross-tissue reproduction and brain negative control

Chromatin-accessible candidate cis-regulatory element (caCRE) calls and per-cluster peak sets were obtained from the Liu et al. 2026 fetal multiomic atlas. Hepatocyte peaks were taken from clusters LI_1, LI_3, LI_4, and LI_6 (29,926 cells, PCW15–22; 44,165 caCREs). Brain peaks were taken from clusters BR_0 through BR_17 (74,035 caCREs). For both tissues, FIMO motif scanning (JASPAR MA0148.1 for FOXA1 and MA0114.2 for HNF4A; p < 10^-3), best-score motif pairing within a 1,001 bp summit-centred window (max inter-motif distance 500 bp), per-bin observed/expected enrichment in 5 bp bins against a 1,000-permutation per-peak null, and Benjamini–Hochberg FDR correction were applied identically to the K562 pipeline. The hepatocyte accessibility comparison (Supplementary Fig. 13) used the HDMA-provided per-peak score (log10(score + 1)); close-spaced (15–60 bp) and far-spaced (100–500 bp) peak groups were compared by one-sided Mann–Whitney U test, with multivariate adjustment for FOXA1 and HNF4A motif counts per peak as a sensitivity analysis.

### Pioneer-seq library design

The Pioneer-seq nucleosome library was based on three nucleosome positioning sequences (NPSs): Widom-601, 5S rDNA, and mouse mammary tumor virus (MMTV)-A^22^. Each NPS was first stripped of pre-existing TF binding sites: FIMO (MEME Suite) and JASPAR 2024 were used to scan each template, and matching bases (FIMO p < 0.01) were modified to remove detected sites. For each TF binding site to be tested — FOXA1 (TGTTTACTTTG, derived from JASPAR MA0148.1), HNF4A (GAGTCCAAAGTCCAG, derived from JASPAR MA0114.2), and nonspecific control (a partial ETS motif, ACCGGAAGTG, derived from JASPAR MA0098.3) — a series of templates was generated containing a single binding site placed at each of 182 dyad-distance positions (−85 to +96 bp relative to the dyad) across all three NPSs (Fig. 4a). For the cobinding library (Fig. 5), an additional series was generated containing one FOXA1 site and one HNF4A site placed adjacently with a fixed 5 bp gap, at each of 77 dyad-distance positions on the same three NPSs. All library sequences were flanked by primer sequences yielding 230 bp total length. The combined library was synthesised as a custom Agilent oligonucleotide pool.

### Nucleosome reconstitution

The Agilent oligonucleotide pool was amplified in 15 PCR cycles using Herculase II fusion DNA polymerase in 100 µL reactions (1× Herculase II reaction buffer, 1 mM dNTPs, 200 pM Agilent library, 250 nM forward and reverse primers). Pooled DNA from 11 reactions was purified (QIAquick PCR purification kit, Qiagen 28104), quantified by NanoDrop, and fragment size was confirmed on a 2% agarose gel. Nucleosomes were generated as previously described^33^ Briefly, nucleosomes were reconstituted by incubating H2A/H2B dimers and H3.1/H4 tetramers (New England Biolabs) with library DNA at an octamer:DNA molar ratio of 1:1.2 in 10 mM DTT, 1.8 M NaCl for 30 min at room temperature, followed by stepwise dialysis (Slide-A-Lyzer MINI 10,000 MWCO; Thermo Fisher 69750) at 4 °C: 1.0 M NaCl 2 h, 0.8 M NaCl 2 h, 0.6 M NaCl 2 h, then TE pH 8.0 overnight. Reconstituted nucleosomes were transferred to BSA-treated tubes (0.3 mg/ml), confirmed by 4% native PAGE, and free DNA was removed by 7%–20% sucrose gradient. Purified nucleosome library was quantified by qPCR and stored at 4 °C for up to one month.

### Nucleosome-binding gel-shift assay

Recombinant human full-length FOXA1 (Origene TP306045) and HNF4A (Origene TP317863) were used for binding reactions. Protein purity was confirmed by Coomassie staining, and binding activity was validated by EMSA using cognate motifs on naked DNA. Binding reactions contained 30 nM purified nucleosome library and matched concentrations of FOXA1 and HNF4A at 0, 15, 30, 60, 120, and 240 nM in DNA-binding buffer (10 mM Tris–HCl pH 7.5, 50 mM NaCl, 1 mM DTT, 0.25 mg/ml BSA, 2 mM MgCl₂, 0.025% Nonidet P-40, 5% glycerol). Reactions were incubated 10 min on ice followed by 30 min at room temperature, then resolved on 4% native polyacrylamide gels (acrylamide:bisacrylamide 29:1) in 0.5× TBE at 100 V at 4 °C. For single-TF Pioneer-seq experiments (Fig. 4), only the indicated TF was titrated at the same concentration series. For cobinding experiments (Fig. 5), FOXA1 and HNF4A were co-titrated at matched concentrations. Three biological replicates were performed (Supplementary Fig. 9). Gels were stained with SYBR-green (Lonza) for visualisation and band excision.

### DNA isolation and sequencing library preparation

Visible and corresponding invisible bands were excised from gels and immersed in diffusion buffer (0.5 M ammonium acetate, 10 mM magnesium acetate, 1 mM EDTA pH 8.0, 0.1% SDS) overnight at 50 °C. Diffusion buffer was filtered through glass wool and DNA was purified with a QIAquick gel extraction kit (Qiagen 28704). Illumina libraries were prepared in two PCR steps: an initial amplification (8–12 cycles, number determined per-sample by qPCR) using four offset primer sets to dephase the libraries; followed by indexing with Nextera dual indices (XT N7xx and S5xx). Reactions were cleaned with AMPure XP beads after each step. Libraries were quantified with the Invitrogen Quant-iT dsDNA assay, pooled in equimolar amounts, and sequenced 2 × 150 bp paired-end on an Illumina NextSeq at the University at Buffalo Genomics and Bioinformatics Core.

### Pioneer-seq data processing

Reads were processed by an automated Snakemake^34^ pipeline. Low-quality 3′ ends were trimmed with cutadapt (-q 30). Forward and reverse reads were merged with vsearch^35^ (--fastq_mergepairs --fastq_minovlen 20 --fastq_maxdiffs 2). Primer sequences were removed with cutadapt^36^; merged reads outside the 174–220 nt range were discarded. FASTQ-to-FASTA conversion used FASTX-Toolkit (fastq-to-fasta). Each read was mapped to the library reference with vsearch (--mincols 150 --id 0.985 --top_hits_only). Per-nucleosome read counts were tabulated per gel band, per concentration, per replicate.

### Relative shift calculation

Prior Pioneer-seq studies quantified TF–nucleosome binding as the relative supershift, defined as log₂((S/S_NS) / (U/U_NS)), where S and U are read counts in the shifted and unshifted bands of the TF lane and the subscript NS denotes paired nonspecific-control nucleosomes. With two TFs co-titrated, the supershifted ladder includes multiple TF–nucleosome stoichiometries and a clean shifted-band quantification is no longer well-defined. We therefore quantified binding as the relative shift^37^ , defined as

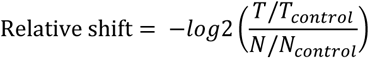

where T and T_control are read counts for the test nucleosome and the paired nonspecific-control nucleosome in the unshifted band of the TF-treated lane, and N and N_control are the corresponding read counts in the unshifted band of the no-TF (null) lane. Nonspecific-control nucleosomes were drawn from the same NPS (Widom-601, 5S, or MMTV) as the test nucleosome and contained only binding sites for unrelated transcription factors, matching nucleosome-positioning properties. The no-TF normalisation controls for sequence-level amplification and capture biases unrelated to TF binding. Depletion of a sequence from the unshifted band in the TF-treated lane, relative to the no-TF baseline and matched nonspecific control, reflects its engagement by the TF. The relative shift is monotonically related to relative supershift for strong binders but is robust to multi-TF complex laddering.

Per-TF binding ability for the single-site library (Fig. 4c) was computed as the mean excess relative shift over the nonspecific control, ⟨RS_TF − RS_NS⟩, averaged across all 182 positions per template. Statistical comparisons between FOXA1 and HNF4A per template used paired Wilcoxon signed-rank tests on the per-position RS_TF − RS_NS values.

### Synergy test on the cobinding library

Per-position synergy on the cobinding library (Fig. 5b) was tested as 2^FH > 2^F + 2^H, where FH is the cobinding RS and F, H are the single-TF RS at the same position and template. Standard errors of 2^x on the linear scale were propagated with the delta method (SE_linear = 2^x · ln 2 · SD_x). The test statistic was a paired z-statistic comparing 2^FH against 2^F + 2^H. Multiple testing across the 77 positions per template was controlled with the Bonferroni correction at α = 0.05; positions with significant synergy are highlighted in Fig. 5b.

### Genomic-nucleosome library and cobinding correlation

Nucleosome dyad positions for the genomic library were inferred from K562 MNase-seq (Mieczkowski et al. 2016; GEO GSM2083140) using the DANPOS algorithm^38^. For each DANPOS-called nucleosome, the center position was expanded to a 191 bp window and scored by a nucleosome-scoring function; the probability assigned to the central base served as the nucleosome-occupancy score, and nucleosomes with a score <0.7 were excluded. Retained nucleosomes that fell within Cooperative-category peaks were scanned for FOXA1 and HNF4A motifs using FIMO (p < 10⁻³, JASPAR MA0148.4 for FOXA1 and MA0114.4 for HNF4A), and only nucleosomes containing exactly one FOXA1 and one HNF4A motif were retained (n = 179). Genomic sequences of the selected 191 bp nucleosome windows were synthesised as described above and reconstituted into the Pioneer-seq library alongside the defined-template constructs. Each genomic nucleosome was assayed under co-titration of FOXA1 and HNF4A, and its relative shift was computed as described. Motif positions relative to the DANPOS-inferred dyad were extracted, and per-nucleosome RS was correlated separately with the FOXA1 motif’s dyad distance and the HNF4A motif’s dyad distance using Pearson and Spearman correlations (computed in R). The two correlations were compared by Fisher’s z transformation (two-sided, independent-samples form; we note this is slightly conservative given both correlations use the same 179 nucleosomes).

### Statistics

Two-sample comparisons used the two-sided Mann–Whitney U test unless stated. Multi-group comparisons used the Kruskal–Wallis test with pairwise Mann–Whitney follow-ups (Bonferroni-corrected). Position-matched comparisons used the paired Wilcoxon signed-rank test. Correlations were Pearson and Spearman; comparison between two correlations used Fisher’s z. Multiple-comparison correction used Benjamini–Hochberg FDR (q < 0.05) unless Bonferroni was specified. All statistical tests were two-sided.

### Software versions

Analyses used Python 3.11 (motif-grammar pipeline) and Python 3.8 (deep-learning pipeline; archived at https://github.com/pwilson97/FOXA1-hnf4a-binding-cnn), with PyTorch 2.2.2, Captum 0.7.0, modisco-lite 2.4.0, scikit-learn 1.3.2, statsmodels 0.14.1, pyBigWig 0.3.22, pybedtools 0.12.0, deepTools 3.5.5, MEME Suite 5.5.7 (FIMO, SpaMo, STREME), bedtools 2.30.0, and samtools 1.23. The Pioneer-seq sequencing pipeline used Snakemake with cutadapt 3.5, vsearch 2.8.1, and FASTX-Toolkit 0.0.14, as described in Wilson et al. (2025) and archived at https://github.com/pwilson97/Pioneer-seq_snakemake. JASPAR 2024 was used for all motif PWMs. Genome assembly: hg19 (GRCh37).

## Data availability

Pioneer-seq sequencing data generated in this study have been deposited at the NCBI Sequence Read Archive under BioProject PRJNA1466564. Re-analyzed public datasets are available at NCBI GEO under accessions GSE182189, GSE182188, and GSE182190 (Hansen et al., 2022a) and GSE104247 (HepG2 ChIP-seq), and at NCBI SRA BioProject PRJNA1482391 with processed data on Zenodo (https://zenodo.org/communities/hdma; Liu et al., 2026). ENCODE K562 chromatin tracks, JASPAR 2024 motif PWMs, and UCSC RepeatMasker hg19 annotations are publicly available from the ENCODE Portal, JASPAR, and UCSC, respectively.

## Notes

### Competing Interest Statement

The authors have declared no competing interest.

### Funding

- National Institute of General Medical Sciences: R35GM158384

