## Supplementary Figures for "Cooperative FOXA1–HNF4A binding emerges from motif spacing and nucleosome architecture"

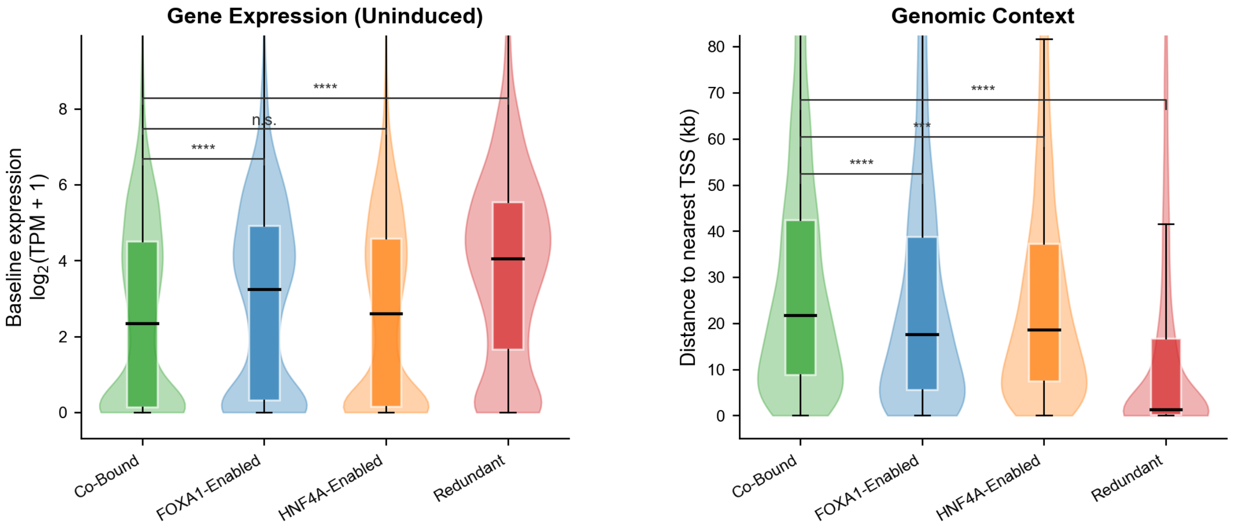


*RNA-seq from Hansen et al. (GSE182190); TSS from UCSC refGene (hg19). Nearest gene within 100 kb. Mann-Whitney U, two-sided. **** p < 0.0001.*

**Supplementary Figure 1 | Baseline gene expression and TSS distance of classified binding sites.** (left) Baseline gene expression at uninduced binding sites. Each classified site (Cooperative, n = 1,824; FOXA1-Enabled, n = 1,510; HNF4A-Enabled, n = 2,727; Redundant, n = 1,875) was assigned to its nearest gene (TSS within 100 kb; UCSC refGene, hg19), and the gene's expression was extracted from RNA-seq of uninduced dual-condition K562 cells (GSE182190; 3 replicates; Salmon quantification). Expression is shown as $\log_2$(TPM + 1), averaged across replicates. Cooperative sites associate with lower-expressed genes (median $\log_2$(TPM + 1) ≈ 2.5) than FOXA1-Enabled sites ($p < 0.0001$) and are not significantly different from HNF4A-Enabled sites. Redundant sites associate with the highest baseline expression ($p < 0.0001$ versus Cooperative). (right) Distance from each binding site summit to the nearest refGene TSS (hg19), in kilobases. Cooperative sites are the most distal from gene promoters (median ≈ 17–21 kb), significantly farther than FOXA1-Enabled ($p < 0.0001$), HNF4A-Enabled ($p < 0.001$), and Redundant sites ($p < 0.0001$). Redundant sites are overwhelmingly promoter-proximal (median ≈ 1–2 kb). In a and b, violins show the underlying data distribution; box plots show median (centre line), interquartile range (box), and 1.5×IQR whiskers. Significance brackets show two-sided Mann–Whitney U tests; **** $p < 0.0001$, *** $p < 0.001$, n.s. not significant.

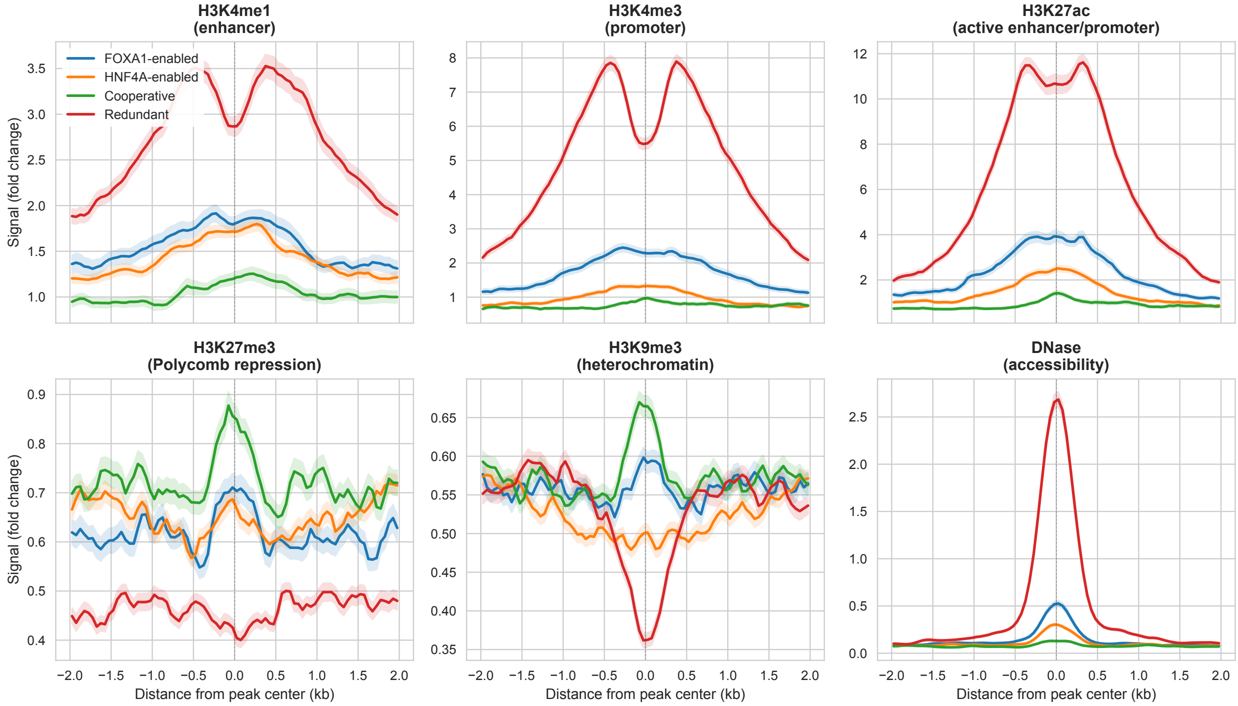


**Supplementary Figure 2 | Histone modification and chromatin accessibility profiles by binding site category.** Metaprofiles of fold-change signal over input across a ±2 kb window centred on peak summits, stratified by binding site category: FOXA1-enabled (FE, blue, n = 1,510), HNF4A-enabled (HE, orange, n = 2,727), Cooperative (CB, green, n = 1,824), and Redundant (red, n = 1,875). Shaded ribbons show 95% confidence intervals across peaks. Top row, active chromatin marks. H3K4me1 (enhancer-associated) is enriched at all four categories, with the highest signal at Redundant sites. H3K4me3 (promoter-associated) and H3K27ac (active enhancer/promoter) are sharply concentrated at Redundant sites, consistent with their promoter-proximal positioning (Supplementary Fig. 1b). Cooperative sites show modest active-mark enrichment. Bottom row, repressive marks and accessibility. H3K27me3 (Polycomb repression) is low across all categories. H3K9me3 (constitutive heterochromatin) is uniformly depleted, indicating that none of the categories reside in constitutive heterochromatin. DNase accessibility is highest at Redundant sites and minimal at Cooperative sites. All histone-modification and DNase tracks are ENCODE K562 baseline (pre-induction). Metaprofiles were computed with deepTools (computeMatrix reference-point --referencePoint center -a 2000 -b 2000 -bs 50).

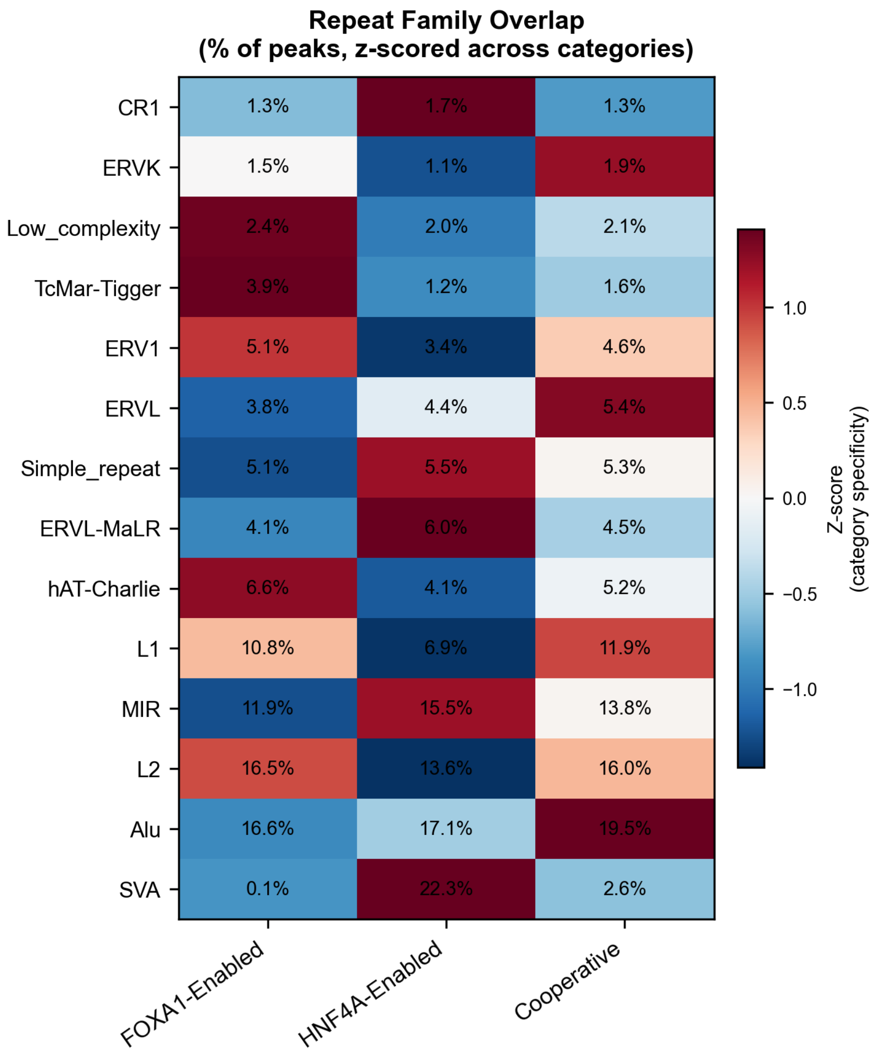


**Supplementary Figure 3 | Repeat family overlap across binding site categories.** Heatmap showing the percentage of peaks in each category overlapping each major repeat family (families with >1% overlap in at least one category). Cell colour encodes category specificity, computed as the z-score across categories within each family (red, enriched relative to other categories; blue, depleted). Raw overlap percentages are annotated in each cell. Repeat families are sorted bottom-to-top in ascending order of their maximum overlap across categories. SVA retroelements (bottom row) are the only family with extreme category-specific enrichment, overlapping 22.8% of HNF4α-Enabled sites versus 0.1% of FOXA1-Enabled, 2.6% of Cooperative, and 3% of Redundant sites. All other families (including Alu, 16.6–19.5%; L2, 13.6–16.5%; L1, 6.9–11.9%; MIR, 11.9–15.5%) show broadly similar overlap rates across categories, consistent with genome-wide background rather than category-specific sequence features. Repeat annotations are from UCSC RepeatMasker (hg19); SVA elements classified as repFamily = "Other" in RepeatMasker were relabelled. Peak coordinates are the classified CUT&Tag narrowPeak files (0.5 μg/ml doxycycline induction).

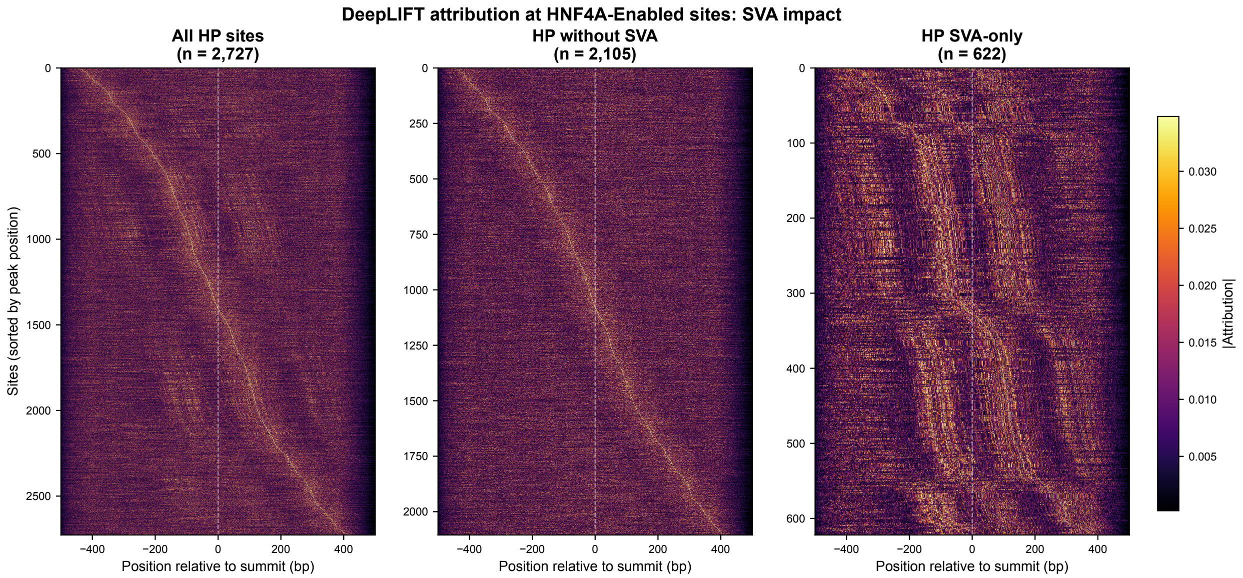


**Supplementary Figure 4 | SVA retroelement overlap produces a split-diagonal artifact in DeepLIFT attribution heatmaps at HNF4α-Enabled sites.** DeepLIFT attribution heatmaps (sum of |attribution| across nucleotide channels) for HNF4α-Enabled (HE) sites, sorted by the position of peak attribution; each row is one site; colour shows per-position |attribution|. Left: all HE sites (n = 2,727) show a bimodal "split diagonal" with two distinct attribution modes separated by ~100 bp. Middle: after removing the 622 sites (22.8%) overlapping SVA retroelements, the pattern resolves to a single clean diagonal (n = 2,105). Right: the 622 SVA-overlapping sites alone show a vertical-stripe barcode pattern corresponding to the fixed positions of DR1-like and forkhead-like sub-sequences within the SVA composite element consensus, confirming that the bimodality reflects stereotyped SVA sequence rather than regulatory grammar. All main-figure attribution and spacing analyses exclude these 622 SVA-overlapping HE sites.

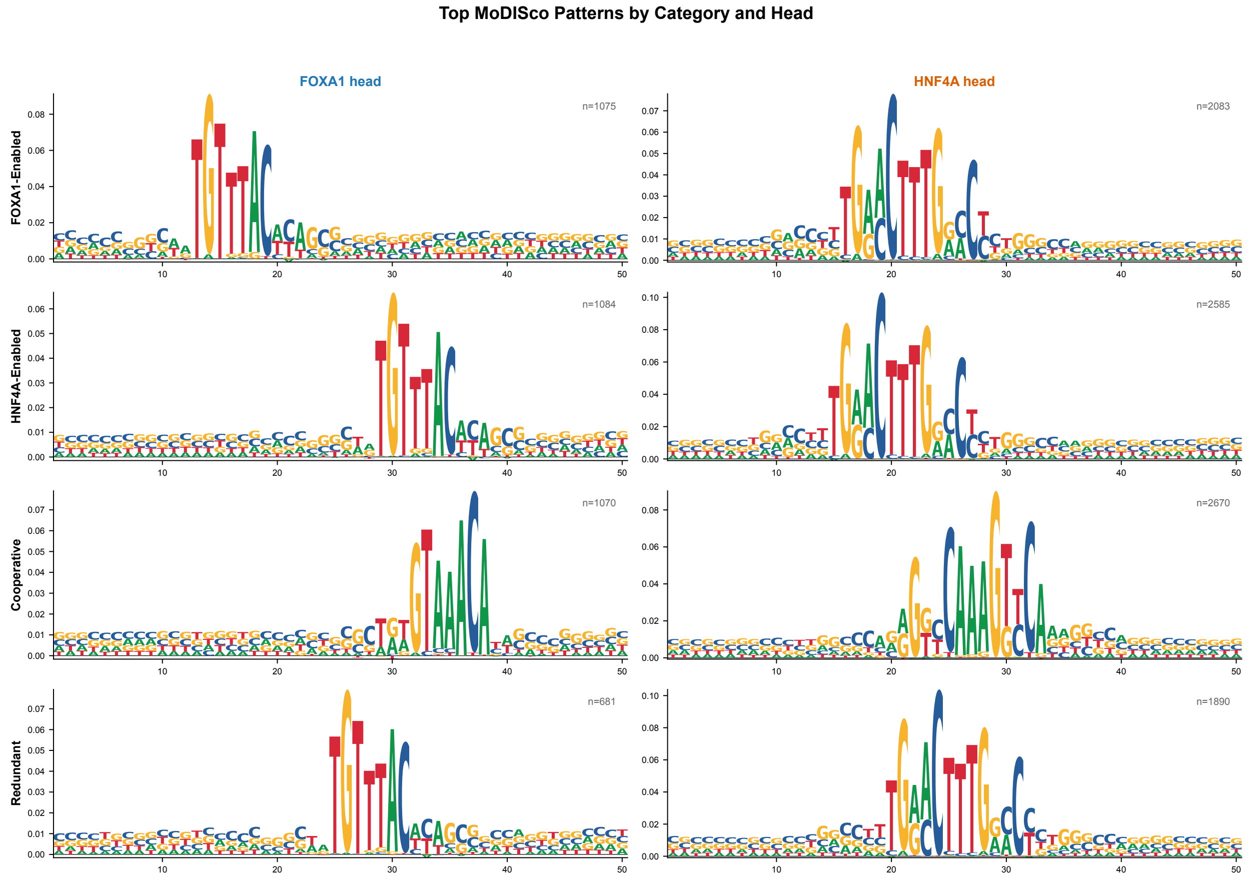


**Supplementary Figure 5 | TF-MoDISco recovers canonical forkhead and nuclear-receptor DR1 motifs from CNN attribution, separately in each site category.** DeepLIFT attribution scores from the dual-head binding CNN on 1,001 bp summit-centred sequences were split by binding site category (rows: FOXA1-Enabled, HNF4α-Enabled, Cooperative, Redundant) and prediction head (columns: FOXA1 head, HNF4α head) for eight independent TF-MoDISco runs (4 categories × 2 heads). Logos show the top-ranked positive contribution-weight matrix (CWM) from each run; letter heights are per-nucleotide attribution contributions, trimmed to informative positions. n per panel is the number of seqlets supporting the CWM. The FOXA1 head recovers a forkhead motif and the HNF4A head recovers a nuclear-receptor direct-repeat-1 (DR1) motif in every category, with no novel or composite motif at Cooperative sites. Motif attribution magnitude is category-dependent: the FOXA1 head reaches its highest per-position attribution at FOXA1-Enabled sites and the HNF4α head at HNF4A-Enabled sites. Attributions were computed on the SVA-filtered dataset (Supplementary Fig. 4).

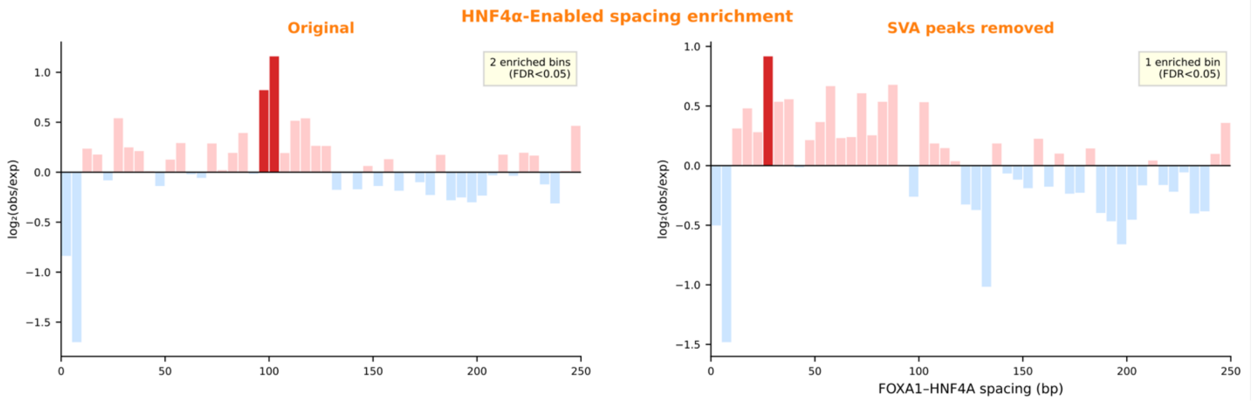


**Supplementary Figure 6 | HNF4α-Enabled spacing enrichment before and after SVA filtering.** FOXA1–HNF4α inter-motif spacing enrichment at HNF4α-Enabled (HE) sites, computed as $\log_2$(observed / expected) in 5 bp bins against a per-peak permutation null (1,000 permutations of motif positions within the 1,001 bp FIMO search window, best-score pairing applied identically). Dark red, bins significantly enriched after Benjamini–Hochberg correction ($q < 0.05$); dark blue, bins significantly depleted; pale bars, not significant. Left: original HE sites (n = 2,727) show 2 enriched bins at ~95–105 bp, the characteristic inter-motif distance within the SVA composite element. Right: after removing the 622 SVA-overlapping peaks (n = 2,105), the ~95–105 bp enrichment is eliminated, leaving 1 enriched bin near 25 bp. The shift in enriched-bin location, away from the SVA-spacing region into the soft-syntax range, confirms that the original ~95–105 bp signal was driven by stereotyped SVA sequence rather than a HE-specific cooperative grammar. Motif pairs identified by FIMO ($p < 10^{-3}$, JASPAR MA0148.1 for FOXA1 and MA0114.2 for HNF4α); best-score pairing strategy; maximum inter-motif distance 500 bp.

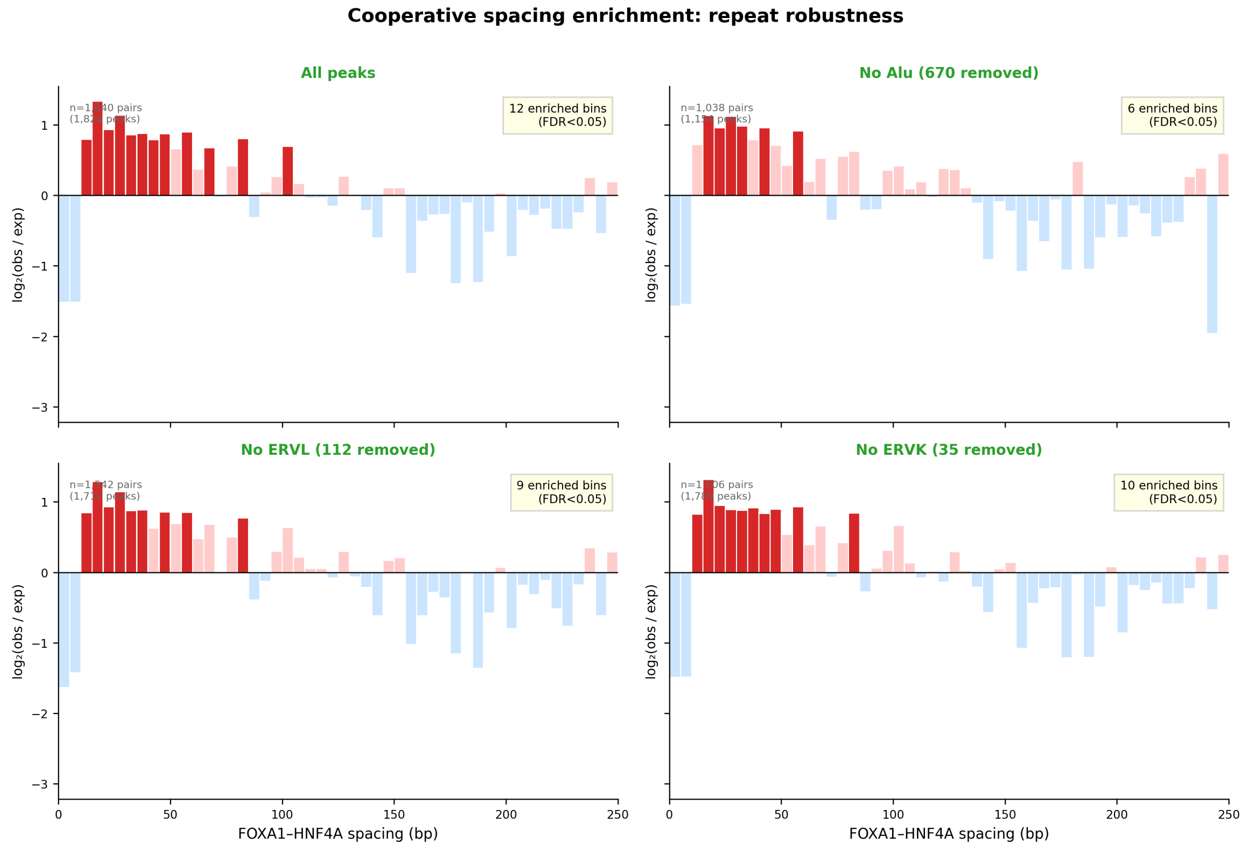


**Supplementary Figure 7 | The Cooperative close-spacing enrichment is robust to removal of CB-enriched repeat families.** $\log_2$(observed / expected) spacing enrichment for Cooperative (CB) sites, computed against a per-peak permutation null (1,000 permutations of motif positions within the 1,001 bp FIMO search window, best-score pairing applied identically) in 5 bp bins. Dark red, bins significantly enriched after Benjamini–Hochberg correction ($q < 0.05$); dark blue, bins significantly depleted; pale, not significant. Repeat families were ranked by their CB category-specificity z-score (computed across categories from the repeat-family enrichment heatmap, Supplementary Fig. 3); the three most CB-enriched families were each tested independently. Top left: all CB peaks ($n = 1{,}824$ peaks; 1,640 motif pairs); 12 enriched bins. Top right: after removing 670 Alu-overlapping peaks (Alu $z = +1.39$, 19.5% of CB peaks; $n = 1{,}154$ peaks; 1,038 pairs); 6 enriched bins. Bottom left: after removing 112 ERVL-overlapping peaks (ERVL $z = +1.30$, 5.4% of CB peaks; $n = 1{,}712$ peaks; 1,542 pairs); 9 enriched bins. Bottom right: after removing 35 ERVK-overlapping peaks (ERVK $z = +1.22$, 1.9% of CB peaks; $n = 1{,}789$ peaks; 1,606 pairs); 10 enriched bins. The 15–60 bp close-spacing enrichment persists after removal of each of the three most CB-enriched repeat families; the reduction in the count of significant bins reflects reduced statistical power from smaller samples rather than loss of signal. This contrasts with HNF4α-Enabled sites, where removal of the 622 SVA-overlapping peaks (SVA $z = +1.41$, 22.8% of HE sites) eliminated the apparent enrichment at ~95–105 bp (Supplementary Fig. 6). Y-axes are shared across all panels.

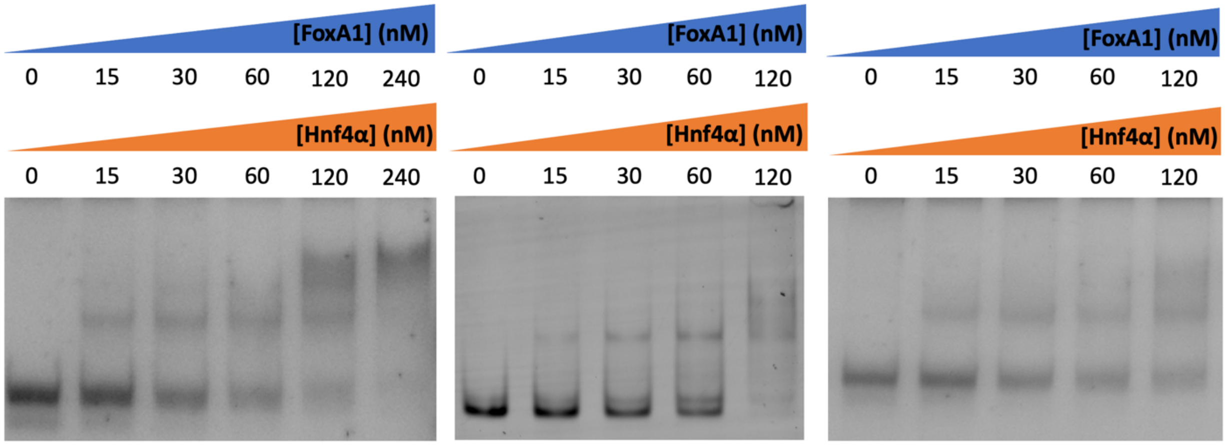


**Supplementary Figure 8 | EMSA titrations of FoxA1+Hnf4α co-titration on the Pioneer-seq nucleosome library across three biological replicates.** Increasing matched molar concentrations of FoxA1 and Hnf4α ([FoxA1] = [Hnf4α]; gradients indicated by blue and orange wedges above each gel) were added simultaneously to 30 nM of the reconstituted Pioneer-seq nucleosome library and resolved on 4% native polyacrylamide gels (acrylamide:bisacrylamide 29:1, 0.5× TBE, 100 V at 4 °C; see Methods) stained with SYBR-green. A: concentration series 0, 15, 30, 60, 120, 240 nM (six lanes). B, C: concentration series 0, 15, 30, 60, 120 nM (five lanes each). On each gel the fastest-migrating (lower) band corresponds to free nucleosome library; intermediate bands correspond to TF–nucleosome complexes of increasing stoichiometry; material near the wells at the highest TF concentrations reflects aggregated or super-shifted species. These gels are the raw input from which the cobinding relative-shift values quantified in Fig. 5 were derived.

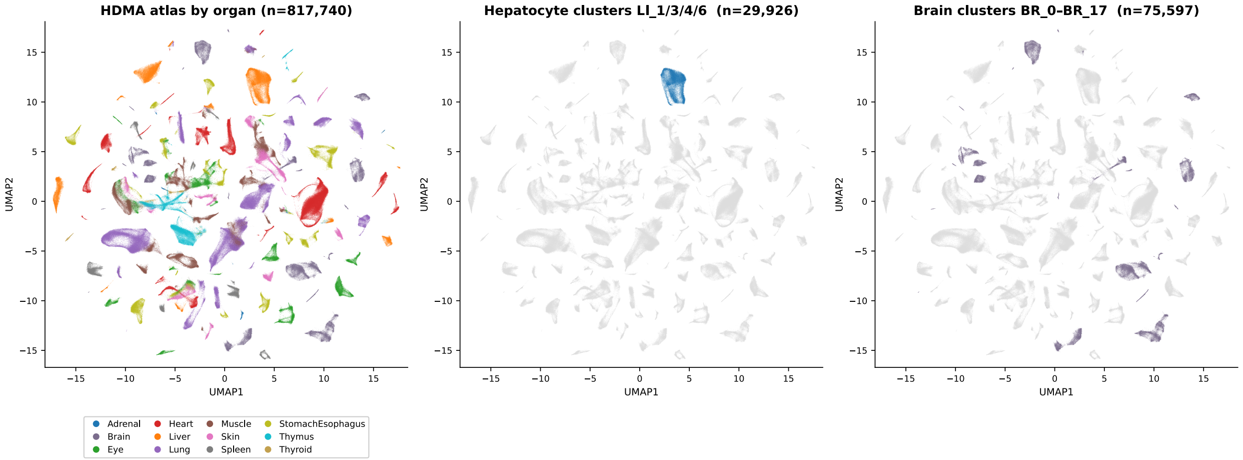


**Supplementary Figure 9 | HDMA single-cell atlas UMAP showing the cell populations used to generate the HDMA Hepatocyte and HDMA Brain panels of main Figure 3C.** UMAP embedding of normalized scRNA expression from the Liu et al. 2026 HDMA fetal multiomic atlas (n = 817,740 cells across 12 organs; BPCells on-disk PCA on the top 3,000 high-variance genes, then uwot UMAP at n_neighbors = 30, min_dist = 0.3, cosine metric). Left: all 817,740 cells colored by organ. Middle: the 29,926 fetal hepatocyte cells from clusters LI_1, LI_3, LI_4, and LI_6 highlighted in blue; caCREs called in these clusters were used for the HDMA Hepatocyte panel of main Figure 3c. Right: the 75,597 fetal brain cells from clusters BR_0 through BR_17 highlighted in grey-purple; caCREs called in these clusters were used for the HDMA Brain (negative control) panel of main Figure 3c.

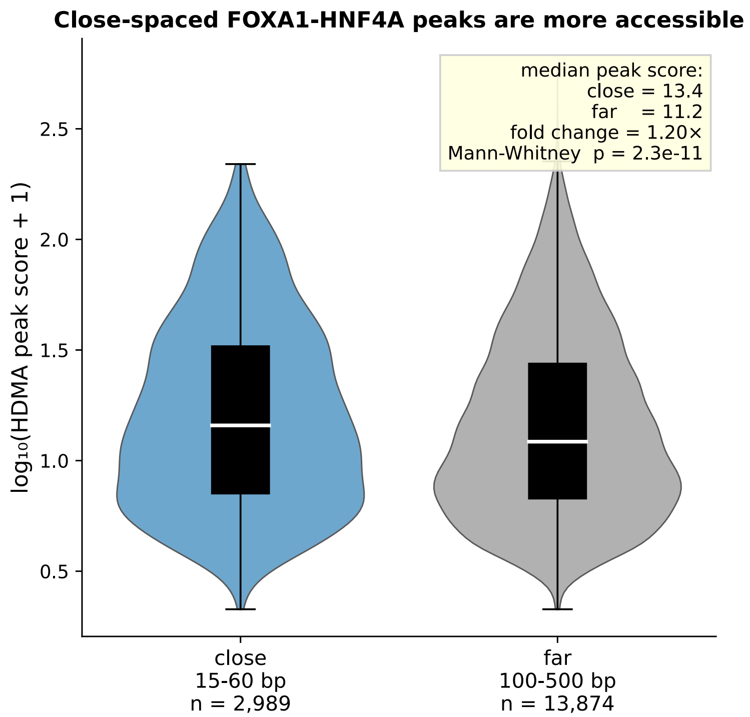


**Supplementary Figure 10. Close-spaced FOXA1-HNF4A peaks are more accessible than far-spaced peaks in HDMA fetal hepatocyte caCREs.**

Per-peak accessibility (HDMA peak score, log10(score + 1)) for HDMA fetal hepatocyte distal caCREs (LI_1, LI_3, LI_4, LI_6; 34,495 peaks; Liu et al., 2026). Both groups contain at least one FOXA1 and one HNF4A motif by FIMO at $p < 0.001$. Close-spaced peaks (n = 2,989) place the best-score FOXA1 to best-score HNF4A center distance within the published 15 to 60 bp soft-syntax window; far-spaced peaks (n = 13,874) place it at 100 to 500 bp. Violins show kernel density; inline boxplots show median (white line) and interquartile range. Median peak score is 13.4 in close-spaced peaks versus 11.2 in far-spaced peaks (1.20-fold; Mann-Whitney U one-sided p = 2.3 x 10^-11). The effect persists after multivariate adjustment for FOXA1 and HNF4A motif counts per peak (1.14-fold, p = 1.4 x 10^-11) and in the most stringent stratum of peaks containing exactly one of each motif (Mann-Whitney p = 0.011). HDMA caCREs are fixed-width (501 bp), so peak length is not a confounder.

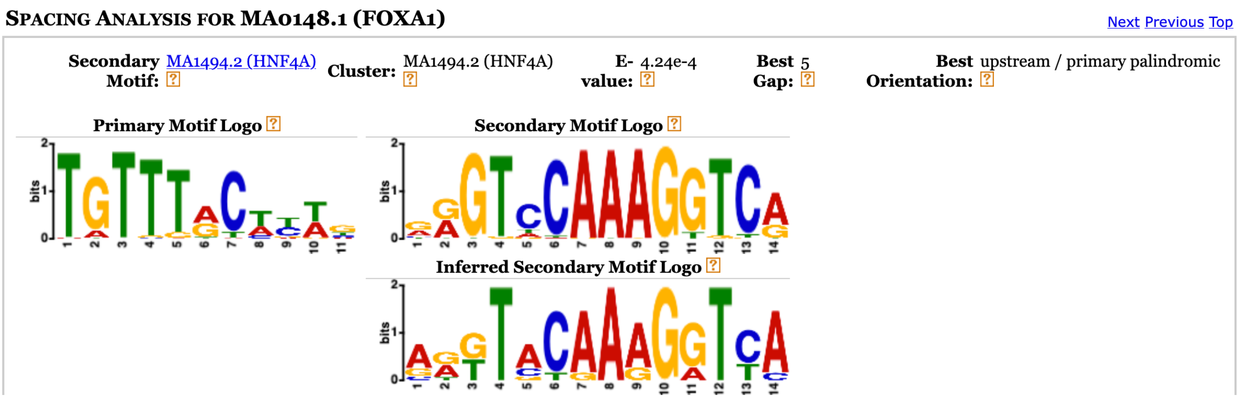


**Supplementary Figure 11 | SpaMo identifies HNF4α as the top-scoring secondary motif at 5 bp upstream of FOXA1 at Cooperative peak sequences.** SpaMo (MEME Suite v5.5.7) was run on Cooperative-category peak sequences (summit ± 500 bp) with the FOXA1 PWM (JASPAR MA0148.1) as the primary motif, against the JASPAR 2024 vertebrate collection. The top-scoring secondary motif was HNF4α (MA1494.2; an alternative JASPAR HNF4α profile to the MA0114.2 used in our FIMO analysis) at a best gap of 5 bp upstream in primary-palindromic orientation (E = $4.24 \times 10^{-4}$). Left: primary motif logo (FOXA1, MA0148.1). Top right: database secondary motif logo (HNF4α, MA1494.2). Bottom right: SpaMo's inferred secondary logo from observed occurrences at the best gap. The inferred and database logos match, confirming the canonical HNF4α DR1 motif.
